# Apcdd1 is a dual BMP/Wnt inhibitor in the developing nervous system and skin

**DOI:** 10.1101/843714

**Authors:** Alin Vonica, Neha Bhat, Keith Phan, Jinbai Guo, Lăcrimioara Iancu, Jessica A. Weber, Amir Karger, John W. Cain, Etienne C. E. Wang, Gina M. DeStefano, Anne H. O’Donnell-Luria, Angela M. Christiano, Bruce Riley, Samantha J. Butler, Victor Luria

## Abstract

Animal development and homeostasis depend on precise temporal and spatial intercellular signaling. Components shared between signaling pathways, generally thought to decrease specificity, paradoxically can also provide a solution to pathway coordination. Here we show that the Bone Morphogenetic Protein (BMP) and Wnt signaling pathways share Apcdd1 as a common inhibitor and that Apcdd1 is a taxon-restricted gene with novel domains and signaling functions. Previously, we showed that Apcdd1 inhibits Wnt signaling, here we find that Apcdd1 potently inhibits BMP signaling in body axis formation and neural differentiation in chicken, frog, zebrafish, and humans. Our results from experiments and modeling suggest that Apcdd1 may coordinate the outputs of two signaling pathways central to animal development and human disease.

**Significance Statement:** *Apcdd1* is a taxon-restricted gene that inhibits both BMP and Wnt intercellular signaling pathways in multiple organisms including mice, frog, zebrafish, and chicken. It encodes a bi-functional protein with a novel protein domain that can bind to Wnt and BMP receptors and block downstream signaling.

## Introduction

Complex multicellular organisms develop through a series of exquisitely choreographed cellular events, such as differentiation and migration (Nusslein-Volhard and Wieschaus, 1980), which must be temporally and spatially coordinated through the synchronized use of signal transduction pathways. Although the specification of cell fate and function by signaling through individual pathways has been well described, the molecular-level intersections of distinct signaling pathways are often unknown.

Less than 20 signaling pathways are used to build and maintain multicellular eukaryotic organisms (Pires-daSilva and Sommer, 2003). Five pathways predominate: Wnt, BMP/Transforming Growth Factor ß (TGF-ß), receptor tyrosine-kinases (RTK), hedgehog and Notch. The Wnt and BMP pathways operate early in development to establish the anterior-posterior and dorsal-ventral axes of the body and generate essential neuronal populations. At later stages, BMP and Wnt are essential for maintaining the integrity and organization of many adult tissues including skin and hair (Lee and Tumbar, 2012). While seminal studies have shown that periodic, sequential and precisely coordinated BMP and Wnt signaling is required during the adult mouse hair cycle (Kandyba et al., 2013; Plikus et al., 2011; Plikus et al., 2008), the molecular nature of pathway coordination and the control of pathway activation dynamics have remained largely unclear.

Signaling specificity and temporal coordination appear to be conflicting requirements: signal specific responses require the regulation of independent components, while temporally coordinating the activation of signaling pathways requires at least some common components. This problem can be resolved by the existence of factors that interact with multiple signaling pathways. Although a few Wnt/BMP pathway coordinators, including Cerberus and Dkk, have previously been identified (Cruciat and Niehrs, 2013; Piccolo et al., 1999), to elucidate the molecular nature of further pathway coordinators, we examined *APCDD1*, a gene mutated in a human hair and skin condition called Hereditary Hypotrichosis Simplex (Shimomura et al., 2010). First, *Apcdd1* has been shown to be induced by the Wnt pathway on tissues that include the skin, retina, central nervous system (CNS) and vasculature (Chen et al., 2015; Daneman et al., 2010; Kuraku et al., 2005; Mazzoni et al., 2017). APCDD1 then inhibits Wnt signaling in both multiple cell types, including the skin, nervous system, and head structures, and multiple species (human, chicken, frog) (Mazzoni et al., 2017; Shimomura et al., 2010). Second, we previously found that *Apcdd1* encodes a dimeric transmembrane protein that complexes with the LRP5 Wnt receptor and lowers Wnt signaling output after activation by the Wnt3a ligand (Shimomura et al., 2010). Third, we noted that human *Apcdd1* heterozygous dominant mutations show more severe skin and hair phenotypes than mouse homozygous loss-of-function mutations (Mazzoni et al., 2017; Shimomura et al., 2010), suggesting APCDD1 may block an additional signaling pathway. Since Wnt depletion also affects BMP targets in various tissues (Hikasa and Sokol, 2013; Lim and Nusse, 2013; Xu et al., 2011), we have assessed the hypothesis that *Apcdd1* is a multi-functional inhibitor of both the BMP and Wnt pathways.

In this paper, we have examined the effect of APCDD1 on BMP signaling during critical stages of cellular patterning during embryonic development. We found that that Apcdd1 is a potent inhibitor of BMP signaling. We find that in addition to its role in modulating WNT signaling, *Apcdd1* directly tunes BMP pathway signaling during gastrulation and tissue patterning, and has a critical role in acquisition of neuronal cell fates. APCDD1 blocks BMP signaling by binding to the BmprIa receptor and inhibiting the nuclear localization of Smad1, the main BMP effector. Using frog, chicken and zebrafish embryos, we find that APCDD1 represses the expression of BMP targets and, interferes with major developmental processes including gastrulation, body axis formation, neural specification and axon pathfinding. APCDD1 is also a Wnt/BMP pathway intersection point: single-cell imaging of BMP and Wnt effectors, coupled with mathematical modeling, show that APCDD1 can coordinate Wnt/BMP activation, suggesting a dynamic explanation for Wnt/BMP periodic sequential activation in mouse hair cycle (Plikus et al., 2011). Finally, *Apcdd1* is an ancient orphan gene that may have arisen *de novo*. It is present in few Metazoan phyla in which BMP signaling and Wnt signaling are used for axial specification, and contains two novel protein domains which resemble each other and are unique to APCDD1 proteins.

## Results

### Apcdd1 inhibits both Wnt and BMP signaling both *in vitro*, and *in vivo* during *Xenopus* embryogenesis

As a first approach to assessing whether Apcdd1 enables the temporal coordination of BMP and Wnt pathway activation, we tested the effect of APCDD1 expression on activation of BMP and Wnt pathways in multiple independent assay systems. First, we measured the levels of phosphorylated Smad1 (pSmad1, activated by BMP signaling) and β-catenin (activated by Wnt signaling) in cultured NIH 3T3 mouse fibroblast cells after BMP and/or Wnt stimulation. Using quantitative immunofluorescence, we found that the levels of pSmad1 and β-catenin were elevated in the nuclei of NIH 3T3 cells by their cognate ligands BMP2 and Wnt3a (Fig. 1A-C). In contrast, pSmad1 and β-catenin were present at low levels in the absence of ligands (data not shown, Fig. 1C). Similarly, the levels of pSmad1 and β-catenin were reduced in cells transfected with *APCDD1* compared to untransfected neighboring cells (Fig. 1A-B, quantification in Fig. 1C), suggesting that Apcdd1 represses the output of both the Wnt and BMP pathways.

**Figure 1.**
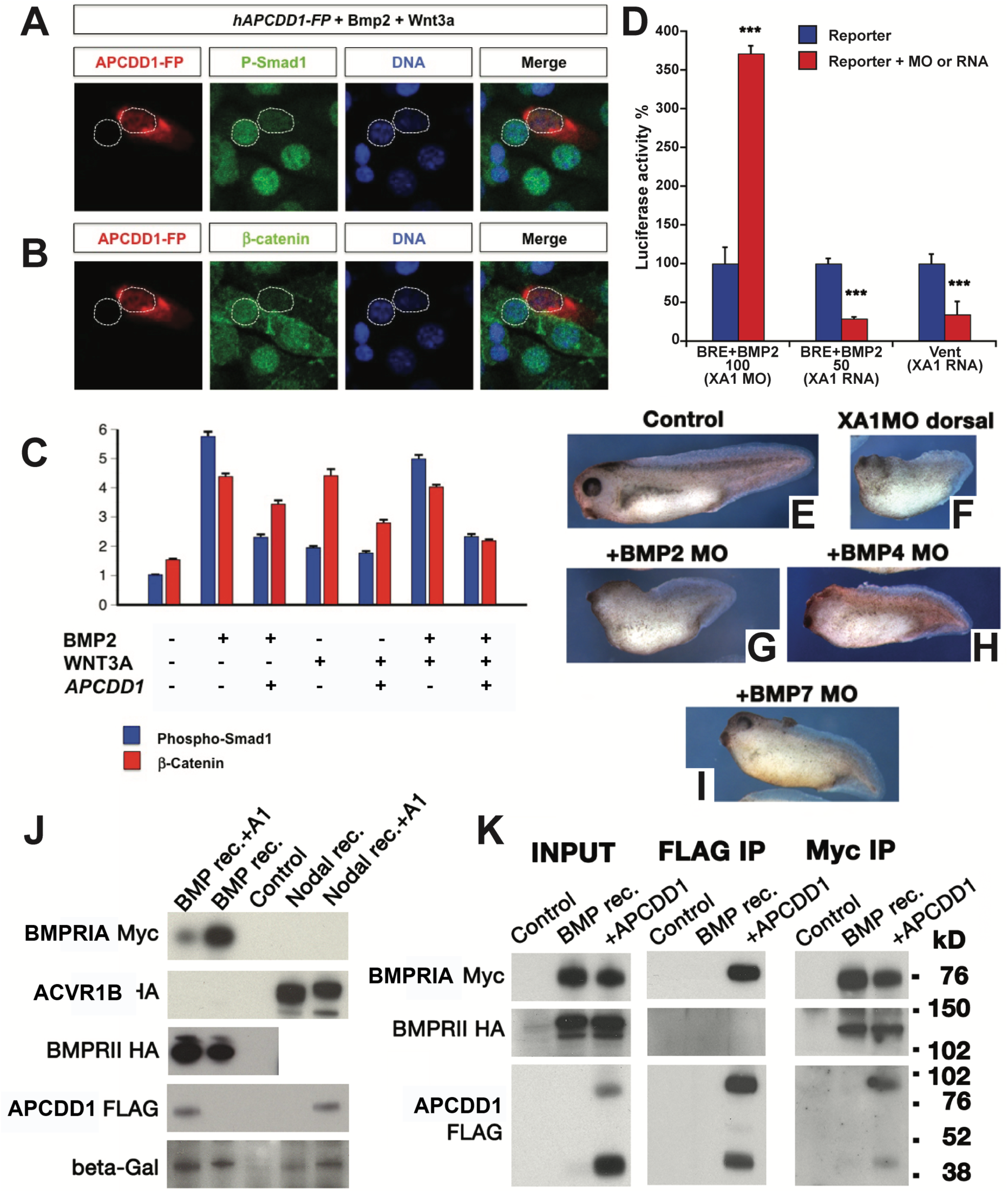
APCDD1 inhibits BMP signaling in cells and tissues. **A-B.** The activation of BMP and Wnt pathways was measured at single-cell level, using NIH 3T3 mouse fibroblast cells cultured in the presence of BMP and Wnt ligands. Cells transfected with *APCDD1* and a fluorescence reporter (displayed in red) have low levels of both pSmad1 (green in A) and β-catenin (green in B), while neighboring cells that are not transfected have higher levels of both effectors. Nuclei are visualized with DAPI (blue). **C.** The immunofluorescence levels of pSmad1 and β-catenin were measured relative to the background, in the presence (0) or absence (1) of one or both ligands (Bmp2, Wnt3A) or of *Apcdd1* (n = 40 cells per condition). The outputs of BMP and Wnt signaling are inhibited by *Apcdd1* (p < 0.001 in Bonferroni-adjusted 2-way heteroscedastic t-test comparisons). **D.** Effect of *Xenopus* Apcdd1 (XA1) modulation on BMP-induced transcriptional activity in *Xenopus* animal injections. Apcdd1 depletion increases transcription induced by exogenous *BMP2* RNA, while overexpression of *apcdd1* RNA reduces transcription from both the BRE reporter induced by exogenous BMP2, and of the Vent promoter induced by endogenous BMP ligands. *** p < 0.001 in Student’s t-test. **E-I.** The phenotype of *XA1* depletion on the dorsal side is rescued most effectively by simultaneous BMP7 depletion (quantification in Fig. S3E). Embryos injected at the 4-cell stage on the dorsal side were collected at stage 36. The dorso-anterior index (DAI) values (Kao and Elinson, 1988) are: E – DAI5 (normal), F – DAI2, G – DAI2, H – DAI3, I – DAI4. **J.** Human APCDD1 specifically reduces the protein level of type I BMP receptor BMPRIA, but not the type II receptor BMPR2 and nodal type I receptor ACVR1B. Beta-galactosidase serves as loading control. *Xenopus* embryos were injected in the animal poles. **K.** APCDD1 interacts directly with BMPRIA, but not with BMPR2, in CHO cells. APCDD1 and BMPRIA coprecipitated in both APCDD1 (center panel) and BMPRIA (right panel) immunoprecipitation. BMPR2 coprecipitated with BMPRIA (right panel), but not with APCDD1 (center panel).

Next, we tested the effect of Apcdd1 on Wnt and BMP pathway signaling using microarray analysis in *Apcdd1*-depleted Xenopus embryos. To identify *Apcdd1* target genes, and further explore its function across developmental pathways, we examined role of *Apcdd1* in the early development of the African clawed frog (*Xenopus laevis*). *Xenopus Apccd1* (XA1) is expressed at neurula and tadpole stages, with an expression pattern similar to that of mouse embryos (Jukkola et al., 2004). Early expression is maternal and zygotic, from 8-cell to gastrula stages, including in the dorsal lip (Spemann organizer) (Fig S1). We performed a microarray analysis of *XA1* morpholino oligonucleotide (MO)-depleted *Xenopus* dorsal cells (Fig. S3A, B). Consistent with results from mouse 3T3 cells, many of the genes that were up-regulated >2-fold in *Apcdd1*-knockdowns were associated with either the BMP pathway (10 out of 26 protein coding genes, Suppl. Table 1) or the Wnt pathway (2 genes). We thus examined whether *Xenopus* Apcdd1 could regulate BMP signaling either *in vivo* (Fig. S2) or in *in vitro* transcription assays (Fig. 1D). We found that MO-depletion of XA1 in *Xenopus* embryos expands the expression of *sizzled*, a downstream target activated by BMP signaling (Fig. S2G-J). The elevated expression of *sizzled* was reduced by the co-injection of a MO-resistant *Apcdd1* RNA (Fig. S2K, L). In contrast, MO-depletion of XA1 reduced the expression of *Sox2,* a negative target of BMP signaling (Fig S2M-S2R). In the transcription assays, we used a BMP Responsive Element (BRE) reporter (Korchynskyi and ten Dijke, 2002) to characterize the extent of BMP2 stimulation in early *Xenopus* embryos (Fig. 1D). Strikingly, depleting endogenous XA1 protein resulted in a >3-fold increase in BMP2 responsiveness, whereas the overexpression of XA1 repressed the ability of BMP2 to activate either the BRE reporter or the endogenous *Vent2* promoter activated by endogenous BMPs (Fig. 1D) (Hata et al., 2000). This finding was also observed in mouse NIH 3T3 cells; cells transfected with human APCDD1 (hAPCDD1) showed a reduced response to BMP4 responsivity (Fig. S3C). We also transfected cells with the mutant form of hAPCDD1 (hAPCDD1-L9R) present in hereditary hypotrichosis and thought to downregulate APCDD1 protein levels by preventing it from reaching the cell membrane (Shimomura et al., 2010). hAPCDD1-L9R was considerably less effective at inhibiting BMP activity in both NIH 3T3 cells (Fig. S3C) and *Xenopus* embryos (Fig. S3D), similar to previous findings related to Wnt-dependent transcriptional activity (Shimomura et al., 2010).

*Apcdd1*-depleted *Xenopus* embryos showed a strong ventralized phenotype, consistent with the idea that APCDD1 normally plays a critical role in tuning BMP signaling. To test this idea, we used morpholinos to quantitatively reduce BMP signaling. Indeed, we found that experimentally inhibiting the BMP pathway rescues the APCDD1*-*depletion phenotype in *Xenopus* embryos. Three BMPs, *Bmp2*, *Bmp4*, and *Bmp7* each have overlapping expression with *Apcdd1* in *Xenopus* (Plouhinec et al., 2011). We generated MOs directed against these BMPs and found that they were able to rescue the ventralized phenotype of *Apcdd1*-depleted embryos (Fig. 1E-I, S3E), with the most effective rescue observed after BMP7 depletion (Fig. 1I, S3E). Of these three BMPs, XA1 was also most effective at suppressing the activity of the BMP7 ligand (Fig. S3F). These results confirm that APCDD1 plays a critical role in tuning BMP signaling during development.

Our data show that in addition to its previously known role in inhibiting Wnt signaling, *Apcdd1* also tunes BMP signaling. Therefore, to determine if Apcdd1 inhibits Bmp signaling directly, we investigated the molecular mechanism of action through which Apcdd1 blocks BMP signaling (Mizutani and Bier, 2008; Moustakas and Heldin, 2009; Umulis et al., 2009). BMP/Growth and Differentiation Factor (GDF) ligands activate a heterodimeric complex of type I and type II BMP serine/threonine receptors (Bmprs) (Mueller and Nickel, 2012). BMP binding leads to the phosphorylation of Bmpr-regulated Smads (R-Smads; Smad1/5/8), which then act as transcriptional regulators. We found that the over-expression of *Xenopus Apcdd1* synergized with *chordin*, a canonical BMP inhibitor, and dramatically reduced signaling through a constitutively active form of the type IA Bmpr (BmprIa, also known as ALK3) (Fig. S3G). Misexpression of *Apcdd1* blocked the phosphorylation of endogenous Smad1 in *Xenopus* animal cap cells (Sasai et al., 1995), with similar effectiveness to *chordin* (Fig. S3H), together suggesting that Apcdd1 may block BMP signaling function by directly interacting with BmprIa. The interaction with Apcdd1 and BmprIa may be specific. We co-expressed *hAPCDD1* together with *BMPRIA* (Alk3), *activin receptor type IB* (*ACVR1B, ALK4*) or *BMPRII* in *Xenopus* embryos (Fig. 1J). Of these three receptors, only the levels of BMPRIA was significantly reduced in the presence of hAPCDD1 (compare ALK3 expression levels in lane 1 and 2, Fig. 1J).

Finally, we performed immunoprecipitation studies to investigate whether there is a direct interaction between Apcdd1 and BmprIa (Fig. 1K). Supporting this hypothesis, BMPRIA-myc immunoprecipitated both APCDD1-Flag and BMPRII-HA in CHO cells (Myc-IP, Fig. 1K). The interaction between APCDD1 and BMPRIA may be specific: while APCDD1-Flag immunoprecipitated BMPRIA-myc, it did not immunoprecipitate BMPRII-HA (FLAG-IP, Fig. 1K). Interestingly, while most APCDD1 protein is monomeric (input, Fig. 1K), the immunoprecipitated form is dimeric (Fig. 1K), suggesting that APCDD1 interacts with BMPRIA as a dimer. Finally, we visualized the BMPRIA-APCDD1 interaction by transfecting CHO cells with both BMPRIA and an inducible form of APCDD1. After APCDD1 induction, BMPRIA appeared to be displaced from the membrane (Fig. S3I) to the perinuclear region (Fig. S3I-L), suggesting that APCDD1 interferes with both the stability and membrane localization of BMPRIA.

### *Apcdd1* is a BMP antagonist in zebrafish

To determine whether the role of *Apcdd1* is conserved between species and at different developmental stages, we examined the function of *apcdd1-like* (*apcdd1l)* during zebrafish embryogenesis. RT-PCR and whole-mount RNA *in situ* hybridization revealed that *apcdd1l* transcript is maternally deposited and is initially found in all cells in the blastula (Fig. S4A). Expression levels subsequently decline during early gastrulation but then increase specifically in dorsal tissues near the end of gastrulation.

Injection of translation-blocking morpholino to knockdown both maternal and zygotic function (Fig. 2A) caused a unique and highly reproducible phenotype characterized by dynamic changes in dorsoventral patterning (Fig. 2). Surprisingly, *apcdd1l* morphants initially appeared dorsalized: expression domains of early organizer genes such as *chd*, *gsc* and *dkk-1* showed marked expansion by 6 hours post-fertilization (hpf; early gastrulation), whereas the domain of ventrolateral marker *vent* showed corresponding reduction at 6 hpf (Fig. 2B, Fig. S4B-D). Because *Apcdd1* also acts as a Wnt-antagonist in diverse vertebrate species (Shimomura et al., 2010), the dorsalized phenotype observed here likely reflects disruption in the role of Apcdd1l restricting maternal Wnt8, a key determinant of the zebrafish organizer (Lu et al., 2011). Evidence of dorsalization persisted until 10 hpf (the end of gastrulation), as shown by expanded domains of *brachyury/ta* in the notochord (Fig. S2E) and *sox19b* and *otx1b* in the neurectoderm (Fig. 2C, S2F), and reduced expression of *tp63* in the ventral/epidermal ectoderm (Fig. S4H). Remarkably, the degree of dorsalization showed a dramatic reversal by 12 hpf (6 somites stage) as revealed by nearly normal patterns of these same regional markers (Fig. 2D, and data not shown). Moreover, by 24 hpf embryos took on a mildly ventralized phenotype in the trunk and tail region as shown by expanded domains of *gata3* and *gata1* in ventral skin ionocytes and hematopoietic tissue, respectively (Hartnett et al., 2010; Hsiao et al., 2007; Lin et al., 2006; Soza-Ried et al., 2010) (Fig. 2G, H).

**Figure 2.**
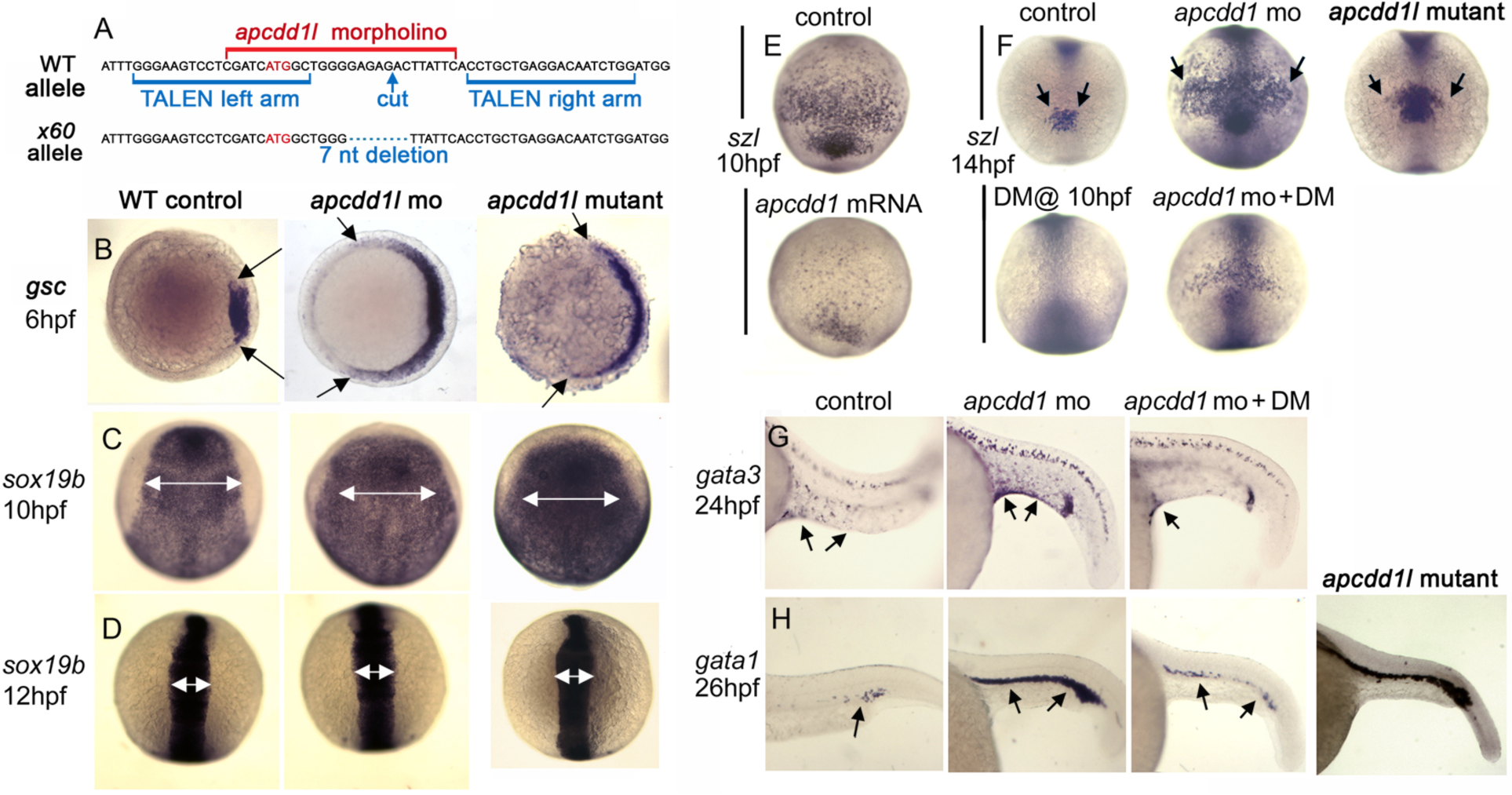
Role of *APCDD1* in zebrafish axis elongation and dorso-ventral patterning. **A.** Partial sequence of exon 1 surrounding the initiation codon (red font) showing the binding sites for *apcdd1l-*mo (red font) and TALEN left and right arms (blue font) used to induce double strand breaks. The induced *x60* mutant allele has a 7-nucleotide deletion, leading to a frame shift followed by 18 premature stops in exon 1 (a presumptive null). **B.** Expression of the organizer gene *goosecoid* (*gsc*) at 6 hpf (early gastrula stage). Knockdown of *apcdd1l*, or loss of *apcdd1l* in MZmutants, disinhibits maternal Wnt, resulting in expansion of the organizer. Animal pole views with dorsal to the right. **C, D.** Expression of neural plate marker *sox19b* at the end of gastrulation (C) and at the 6-somite stage (D). The early expansion of *gsc* in *apcdd1-*morpants and MZmutants results in a dorsalized phenotype that persists through the end of gastrulation, but dorsalization is rapidly reversed by early somitogenesis stages. Dorsal views with anterior to the top. **E.** Expression of the Bmp feedback inhibitor *sizzled* (*szl*) in ventral ectoderm (anterior up) at the end of gastrulation. Expression of *szl* marks cells experiencing active Bmp signaling. Injection of *apcdd1l* mRNA into wild-type embryos at the one-cell stage results in downregulation of *szl* by the end of gastrulation. **F.** Expression of *szl* in ventral ectoderm (anterior up) at the 10-somite stage. *apcdd1l-*morphants and MZmutants exhibit an expanded ventral domain of *szl*, indicating that Bmp signaling is now higher than normal. Treatment of embryos from 10 hpf with 75 µM dorsomorphin (DM) abolishes *szl* expression in control embryos and partially reverses expansion of *szl* expression in *apcdd1l-*morphants. **G, H.** Expression of *gata3* in ventral epidermis ionocytes (arrows) at 24 hpf (G) and *gata1* in blood progenitors (arrows) at 27 hpf (H). Ventral ionocytes and blood progenitors are expanded in *apcdd1l*-morphants, indicating ventralization of caudal structures. *MZapcdd1l* mutants also show an expanded domain of *gata1*. Treatment of *apcdd1l-*morphants with 75 µM DM from the end of gastrulation partially reverses ventralization. Images show lateral views with anterior to the left.

We hypothesized that the rapid shift from a dorsalized to ventralized phenotype resulted from rising levels of BMP activity near the end of gastrulation due to disruption of Apcdd1l-mediated BMP antagonism. In support of this hypothesis, the expression of *bmp4* was upregulated by the end of gastrulation in *apcdd1l* morphants (Fig. S4G). Additionally, expression of *sizzled* (Salic et al., 1997), a direct feedback inhibitor of BMP signaling (Yabe et al., 2003) began to be upregulated and expand by 8 hpf in *apcdd1l* morphants, several hours before the shift in dorsal-ventral (DV) patterning. *Sizzled* expression remained expanded through at least 14 hpf (10 somites stage) (Fig. 2F). In contrast, injection of wild-type *apcdd1l* mRNA strongly reduced *sizzled* expression by 10 hpf (Fig. 2E). Finally, treating *apcdd1l* morphants at 10 hpf with dorsomorphin (DM), a pharmacological inhibitor of BMP signaling (Kwon et al., 2010; Yu et al., 2008), strongly reduced expression of *sizzled* at 14 hpf (Fig. 2F) and restored normal expression of *gata1* and *gata3* in ventral tissues (Fig. 2G, H). These data support the hypothesis that zebrafish Apcdd1l is required near the end of gastrulation to limit BMP activity.

To validate the morpholino knockdown phenotype, we used TALENs to target exon 1 near the translation start site and recovered a knockout allele (termed *x60*) with a 7-nucleotide deletion (Fig. 2A). The mutant allele leads to a frame shift and a series of premature stops, likely a null. Subsequent breeding showed that the *apcdd1l* mutation is homozygous-viable, enabling the generation of mutant embryos lacking both maternal and zygotic function (MZmutants). MZmutants appeared very similar to *apcdd1l* morphants, showing a markedly dorsalized phenotype during gastrulation (Fig. 2B, C). Dorsalization was later reversed (Fig. 2D) as MZmutants became mildly ventralized (Fig. 2H). Moreover, the ventral domain of *sizzled* expression was enlarged at 10 somites stage relative to wild-type embryos (Fig. 2F), though not to quite the extent of *apcdd1l* morphants. Thus, both mutant and morphant phenotypes support the role of Apcdd1l as a BMP antagonist in zebrafish.

### *Apcdd1* misexpression blocks BMP-dependent activities in the developing spinal cord

To test the ability of Apcdd1 to inhibit BMP signaling in a heterologous system and in later developmental contexts, we examined the effects of *Apcdd1* mis-expression during chick embryogenesis. It has been previously shown that BMP signaling has multiple roles during the development of the vertebrate dorsal spinal cord. First, BMPs and activin act from the roof plate (RP) to direct the fate of dorsal interneurons (dI) 1 to dI3 (Chizhikov and Millen, 2005; Liem et al., 1997) (Fig 3G). The more ventral dorsal neurons form independently of RP signals (Lee et al., 2000; Timmer et al., 2002; Wine-Lee et al., 2004). Blocking BMP signaling using the inhibitory (I) Smad, Smad7, results in the loss of dI1 and dI3 (Hazen et al., 2011). Second, BMP signaling controls the rate of dI1 (commissural) axon growth (Hazen et al., 2012; Phan et al., 2010; Yamauchi et al., 2013). When BMP signaling is blocked by mis-expression of another I-Smad, Smad6, dI1 axon growth stalls (Hazen et al., 2011). While *Apcdd1* is expressed in the anterior neural tube, it is only present in blood vessels at spinal levels (Fig. S5), making it unlikely that APCDD1 has an endogenous role controlling BMP signaling in the dorsal spinal cord. Nevertheless, the absence of *Apcdd1* expression in the spinal cord provides an opportunity to examine whether APCDD1 has analogous activities to the I-Smads.

**Figure 3.**
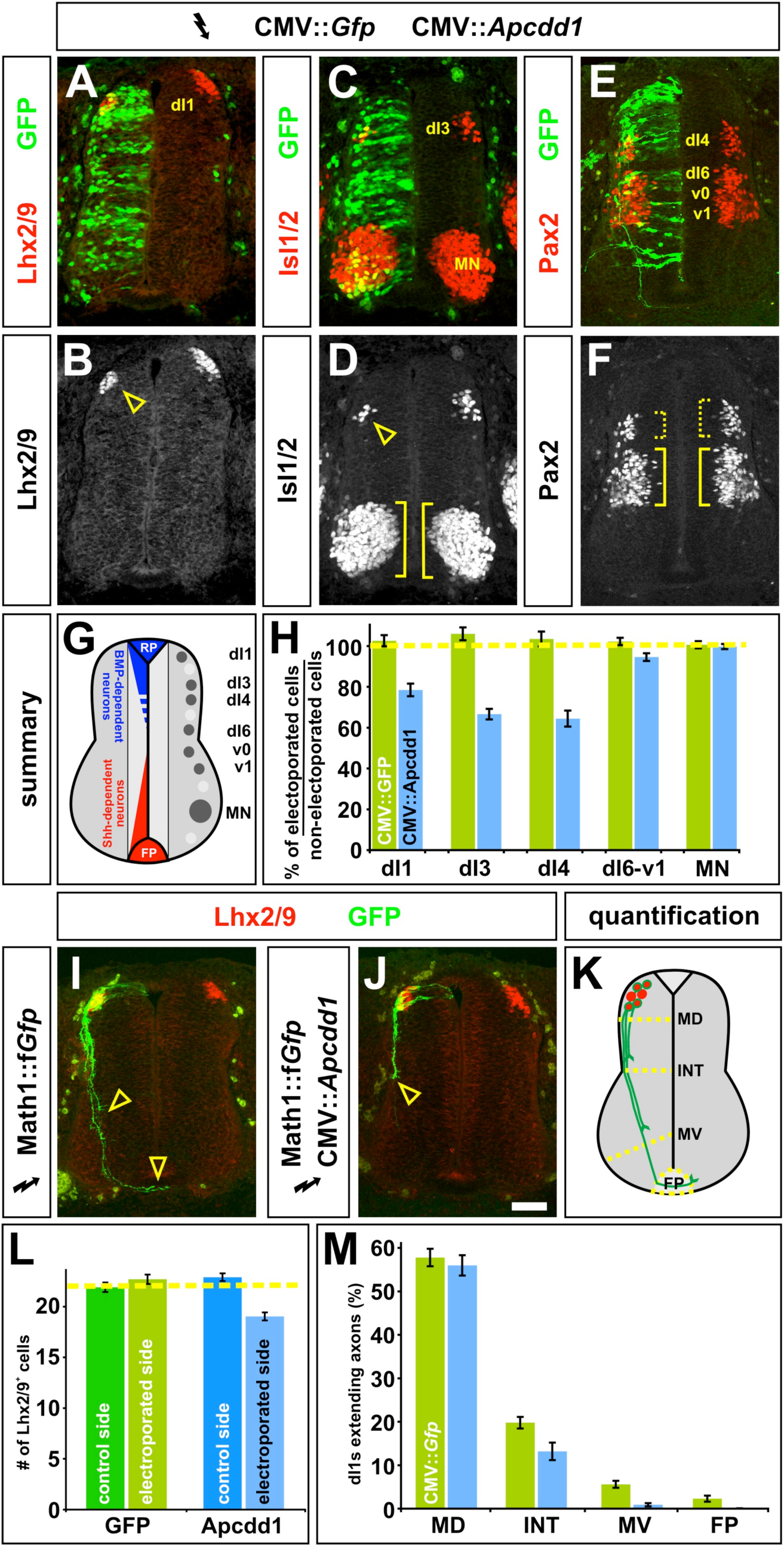
Mis-expression of *Apcdd1* blocks BMP-specific activities in the dorsal spinal cord. **A-F, H.** *Gfp* alone (control, H), or *Gfp* (green) in combination with *Apcdd1* (experimental, A-F, H), was ectopically expressed throughout the spinal cord under the control of the CMV enhancer by *in ovo* electroporation at Hamilton Hamburger (HH) stages 14/15. Embryos were harvested at HH stage 23 and examined for the number of Lhx2/9^+^ dI1 (commissural) neurons (red, A, B), Isl1/2^+^ dI3 and motor neurons (MNs) (red, C, D) or Pax2^+^ dI4 and dI6-v1 neurons (red, E, F). **G.** The relative positions of the labeled post-mitotic neurons are shown schematized within the spinal cord. The dorsal most populations of spinal neurons (dI1-dI3) are known to be dependent on BMP signaling, the role of BMP signaling in the more ventral-dorsal populations (dI4-dI6) has remained unresolved. **H.** Quantification revealed that the misexpression of *Gfp* (n=42 sections from 6 embryos) had no significant effect on any of the monitored populations of spinal neurons (dI1, student’s *t*-test p>0.38 compared to the non-electroporated control side of the same embryo; dI3, p>0.19; dI4, p>0.40; dI6-v1, p>0.35; MN, p>0.49). In contrast, the ectopic expression of both *Gfp* and *Apcdd1* (n=35 sections from 6 embryos) resulted in a 25-35% loss in BMP-dependent dorsal neural populations: dI1 neurons (p<8.1×10^-8^ different from the GFP^+^ control electroporation) dI3 neurons (p<1.8×10^-9^) and dI4 neurons (p<2.3×10^-6^). However, *Apcdd1* had no detectable effect on the numbers of BMP-independent neurons, specifically dI6, vo, v1 interneurons (p>0.12) and MNs (p>0.49). **I.** Lhx2/9^+^ dI1 neurons (red) electroporated with farnesylated (f) *Gfp* under the control of the Math1 (Atoh1) enhancer (Math1::f*Gfp*) at HH stages 16/17, GFP^+^ axons (green) are extending into the ventral spinal cord by HH stage 22, with a few axons having reached the floor plate (FP, arrowheads). **J.** In contrast, Lhx2/9^+^ dI1 neurons (red) concomitantly electroporated with both Math1::f*Gfp* (green) and CMV:: *Apcdd1* showed over an 80% reduction of axon outgrowth at the same stage, with no axons reaching the FP (arrowhead). **K.** The extent of the dI1 axon outgrowth was quantified by determining whether dI1 axons had crossed one of four arbitrary lines in the spinal cord: mid-dorsal (MD), intermediate (INT), mid-ventral (MV) or the FP, as in (Phan et al., 2010). **L.** Misexpression of *Apcdd1* throughout the spinal cord at HH stage 16/17 has a small but significant effect on the numbers of dI1 neurons. There is a 10 % reduction in dI1 neurons on the *Apcdd1* electroporated side (p<2.2×10^-9^, n=173 sections from 6 embryos) compared to the control *Gfp* electroporated side (n=178 sections from 5 embryos). **M.** By HH stage 22, approximately 55 % of Lhx2/9^+^ neurons electroporated with *gfp* (control) or *gfp* with *Apcdd11* (experimental) have extended GFP^+^ axons. In control embryos (n=178 sections, taken from 5 embryos), 10% of these axons extended to the MV line and 4% reached the FP. In contrast, only 1.6% of Lhx2/9^+^ neurons in the embryos electroporated with both *Gfp* and *Apcdd1* (n=173 sections, taken from 6 embryos) had extended to axons to the MV line (p<8.5×10^-8^ similar to GFP controls) with none reaching the FP (p<7.6×10^-4^). Scale bar: 25 μm.

To test this hypothesis, we ubiquitously mis-expressed *Apcdd1* in chicken spinal cords using *in ovo* electroporation. When *Apcdd1* was expressed at high levels immediately before neurogenesis commenced in the dorsal spinal cord (Hamilton Hamburger (HH) stages 10/12), a severe proliferation defect was observed throughout the spinal cord, as previously observed (Shimomura et al., 2010). However, if *Apcdd1 1* was mis-expressed at lower levels while neurogenesis was ongoing (HH 14/15), a different phenotype was observed. There was a 25-35% loss of the dI1 (Fig. 3A, B, H), dI3 (Fig. 3C, D, H) and dI4 (Fig. 3E, F, H) neurons, a similar consequence to Smad7 mis-expression (Hazen et al., 2011). The ectopic expression of *Apcdd1* had no detectable effect on the BMP-independent classes of spinal neurons, strongly suggesting that *Apcdd1* specifically blocks the ability of BMP signaling to induce dorsal cell fates.

We next examined whether the presence of Apcdd1 affected dI1 axon growth by mis-expressing *Apcdd1* at HH stages 16/17, as dI1 axons extend axons away from the dorsal midline (Fig. 3K). The Apcdd1^+^ GFP^+^ dI1 axons displayed significant defects in axon extension (Fig. 3J) compared to GFP^+^ control axons (Fig. 3I). We quantified axon outgrowth by determining whether dI1 axons had crossed one of four arbitrarily drawn lines in the spinal cord: mid-dorsal (MD), intermediate (INT), mid-ventral (MV) spinal cord, or the FP (Fig. 3K). By HH 22, similar numbers of control and experimental dI1s had extended axons (Fig 3M). 10% of control GFP^+^ axons then projected into the ventral spinal cord with 4% reaching the FP. In striking contrast, only 1.6% of Apcdd1^+^ dI1 neurons extended axons to the MV line and none reached the FP, a phenotype similar to that observed after the mis-expression of Smad6 (Hazen et al., 2011).

### *Apcdd1* is expressed in hair follicles

Finally, we characterized the putative role of Apcdd1 as a dual BMP/Wnt inhibitor in mammals. We previously reported the localization of APCDD1 in the dermal papilla (DP), matrix (Mx), hair shaft cortex (HSCx), and cuticle (HSCu) of the human hair follicle (Shimomura et al., 2010). Here, we examined the expression of *Apcdd1* in the skin and hair of adult mice. We found that *Apcdd1* is expressed in the mature keratinocytes of the inner root sheet (Fig. 4A-C), consistent with its role in downregulating both Wnt signaling (Shimomura et al., 2010) and BMP signaling. While previous experiments in mouse revealed coordinated and periodic asynchronous BMP and Wnt signaling activation in mouse hair follicles ((Kandyba et al., 2013; Plikus et al., 2011; Plikus et al., 2008), the molecular nature of the coordination of temporal dynamics remained elusive. Since Apcdd1 is expressed at the right time and place in the hair follicles, we set out to determine if it may control such dynamics. Thus, to gain quantitative insight into the manner in which BMP and Wnt pathways are coordinated in the hair follicle, we developed a model with coupled differential equations, whose variables represent experimentally measurable BMP and Wnt signaling components, including *Apcdd1*. As in similar genetic circuits (Suel et al., 2007), equations represent known interactions between circuit components and use kinetic parameters inferred from our experiments. First, the model reproduces our data and predicts the activation of two BMP and Wnt pathway outputs, pSmad1 and β-catenin (Fig. S6), as a function of BMP and Wnt inputs (Kandyba et al., 2013; Miyoshi et al., 2012; Song et al., 2014). Second, we modelled cases relevant to hair follicle dynamics (Fig. 4D), where BMP and Wnt ligand inputs are constant or periodic, and predict that interesting temporal dynamics may emerge if ligands are periodic, as happens in *in vivo* in hair and skin (Kandyba et al., 2013; Plikus et al., 2011; Plikus et al., 2008). When BMP and Wnt oscillate out of phase, as *in vivo* in mouse hair follicles (Plikus et al., 2011), the presence of Apcdd1 should shorten the time when BMP effector pSmad1 is at high levels (Fig. 4D).

**Figure 4.**
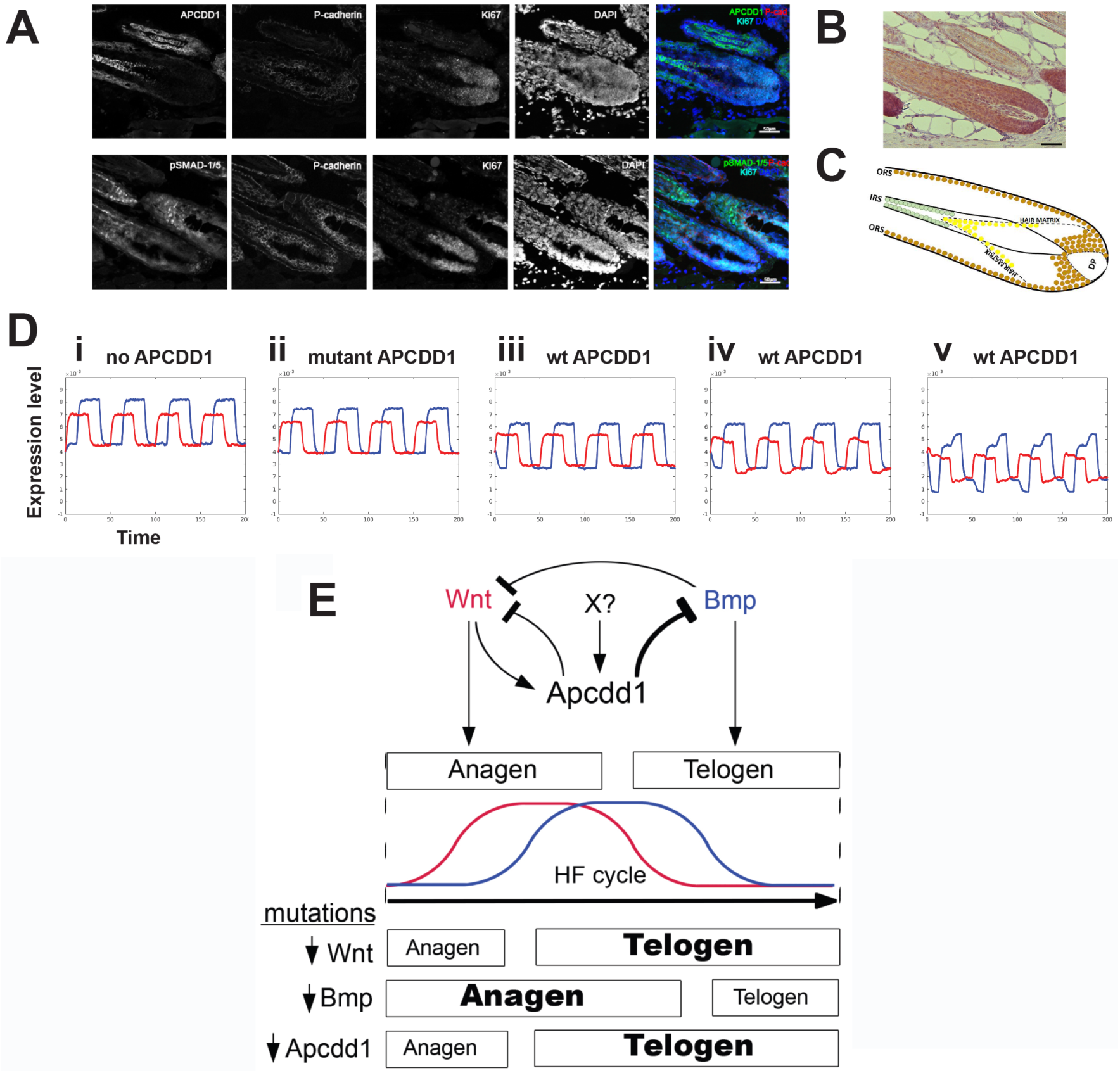
*Apcdd1* is a taxon-restricted gene that encodes a protein expressed in hair follicles and has novel domains. **A.** Immunofluorescence staining of adult mouse hair follicle cryosections detects expression of Apcdd1 (upper row) and respectively pSmad1/5 (lower row) in relation to hair follicle matrix marker P-cadherin and proliferation marker Ki67. The two sections are 20 microns apart. Nuclei are visualized with DAPI. **B.** In mouse hair follicles paraffin sections, activated β-catenin (brown) is present in the hair matrix abutting the dermal papilla (DP) and in the outer root sheet (ORS). Scale bar: 50 μm. **C.** Schematic of immunodetection results. Activated β-catenin (brown), an effector of Wnt signaling, is localized along the outer root sheath (ORS) and hair bulb, surrounding the dermal papilla (DP) that produces the growth signals during anagen. Nuclear pSMAD-1/5 (yellow), an effector of BMP signaling, is located in the distal portion of the hair matrix, where keratinocytes begin to differentiate into the layers of the inner root sheath (IRS). Apcdd1 expression (green) is localized to the mature keratinocytes of the IRS, consistent with its function to downregulate both Wnt and BMP signaling. **D.** Mathematical model of the hair follicle temporal activation dynamics of the BMP and Wnt pathways (pSmad1 in blue, β-catenin in red). The BMP and Wnt ligand inputs are periodic, have the same period and are out of phase by a quarter of a period. Cases shown: (i) no Apcdd1 is present, as in null mutants; (ii) mutant Apcdd1, which is a very weak inhibitor (Shimomura et al., 2010); (iii) wild type Apcdd1, which inhibits Wnt signaling (Shimomura et al., 2010) and BMP signaling; (iv) wild type Apcdd11, and added moderate repression of Wnt signaling by Bmp signaling; (v) wild type Apcdd1, added moderate repression of Wnt signaling by Bmp signaling, and Apcdd1 is induced by Wnt signaling (Fig. 5). The absence of APCDD1 allows higher levels of pSmad1; wild type Apcdd1 depresses pSmad1 levels. **E.** *Apcdd1* controls the duration of the phases of the hair follicle cycle. Summary of Apcdd1 activity: First, *Apcdd1* expression is induced by Wnt signaling and also by a currently unidentified factor X, which maintains Apcdd1 after Wnt signaling ceases. Second, Apcdd1 represses both Bmp signaling and Wnt signaling (which also is repressed by Bmp signaling). Bmp and Wnt signaling together control the duration of hair follicle phases anagen (when hair grows) and telogen (when hair falls out). Third, mutations inactivating Wnt signaling, or Bmp signaling, or Apcdd1, result in changed durations of anagen and telogen.

### *Apcdd1* has an evolutionarily novel protein domain

To understand the origin of *Apcdd1* and find whether it has protein domains, we mapped its evolutionary emergence and analyzed its predicted protein structural features. First, we found *Apcdd1* is present in only 10 of the 35 animal phyla (Fig. 5A). *Apcdd1* is expanded in 2 phyla (Chordates and Arthropods) and is syntenic within, but not between, phyla (data not shown). While *Apcdd1* appears in Metazoans but not in fungi or plants (Fig. 5A), we did find a 205 amino-acid protein resembling human APCDD1 in a few bacterial species. *Apcdd1* is similar to orthologous genes in several other genomes, but to no other genes, suggesting *Apcdd1* is a taxon-restricted gene that may have arisen at the origin of Metazoa. Interestingly, *Apcdd1* was maintained in a few metazoan phyla where it acquired useful functions in Wnt and BMP signaling, which in these phyla control axial specification during embryonic development (Angerer et al., 2000; Darras et al., 2018; Guder et al., 2006; Henry et al., 2010; Henry et al., 2008; Hogvall et al., 2014; Kuo and Weisblat, 2011; Lambert et al., 2016; Lowe et al., 2006; Marlow et al., 2014; Molina et al., 2011; Pang et al., 2011; Pang et al., 2010; Park and Priess, 2003; Plickert et al., 2006; Rentzsch et al., 2007; Schenkelaars et al., 2017; Scimone et al., 2016; Suzuki et al., 1999; Treffkorn and Mayer, 2013; Wikramanayake et al., 1998; Windsor and Leys, 2010) and was lost in phyla that do not utilize Bmp for axial specification have lost APCDD1 (Fig. 5A, Supp. Table 2). Second, we examined the secondary structure content (Yachdav et al., 2014) of 441 Apcdd1 protein sequences (Fig. S7). We found low disorder, and 45% secondary structure elements (16 % α-helix, 29% β-strand). The β-strand content of Apcdd1 proteins is higher than that of ancient proteins (Schaefer et al., 2010) but similar to that of new proteins (Abrusan, 2013). By aligning multiple Apcdd1 sequences, we found that Apcdd1 proteins have two ∼210 amino acid novel domains that resemble each other, are present only in Apcdd1 proteins in different species, and likely arose by internal duplication (Fig. 5B). The Pfam database (El-Gebali et al., 2019) has also recently annotated this domain as Apcddc. Since the signal peptide and the transmembrane domain are not present in all Apcdd1 proteins, Apcdd1 subcellular localization may vary between species. Thus Apcdd1 is a taxon-restricted gene that encodes a protein with a novel protein domain and inhibits BMP and Wnt signaling in organisms that use BMP and Wnt for embryonic axial specification.

**Figure 5.**
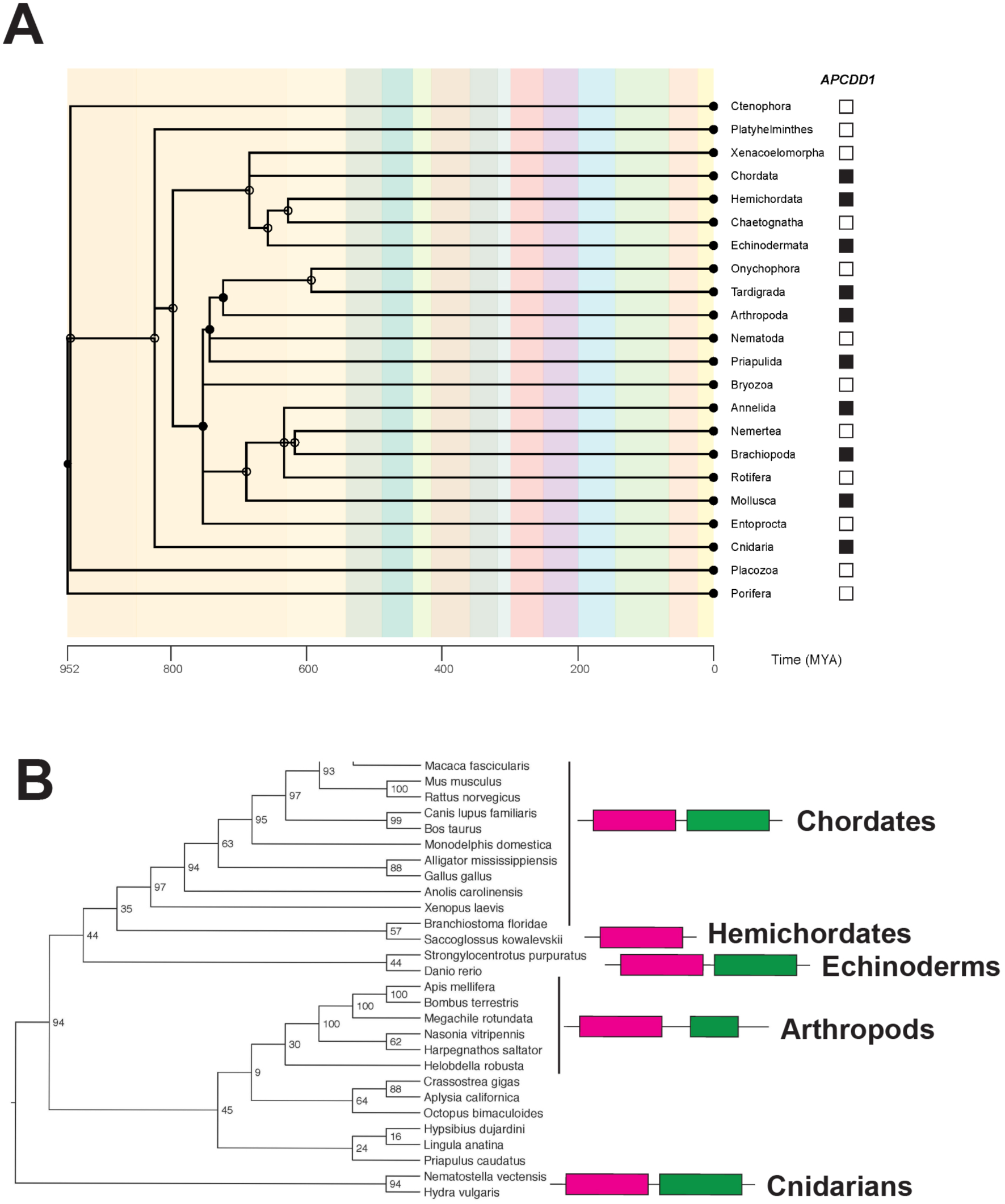
Evolutionary origin of *Apcdd1* and Apcdd1 new protein domains. **A.** Apcdd1 is present in 9 of 35 Metazoan phyla. Species divergence tree in animal phyla having at least one fully sequenced genome. TimeTree.org was used to generate the phylogeny by specifying Metazoa (group) and phylum (rank), and displaying time (MYA) (Kumar et al., 2017). Solid circles denote nodes that mapped directly to the NCBI taxonomy; open circles denote nameless nodes that were not listed as higher-level taxa in the Taxonomy browser. The presence of the *Apcdd1* gene in at least one species of the phylum is indicated by a black box, while the absence of *Apcdd1* is indicated by a white box. The most parsimonious explanation suggests an early Metazoan origin of the Apcdd1 gene, followed by repeated gene loss in multiple phyla. **B.** Apcdd1 protein sequence divergence tree. 30 protein sequences similar to human Apcdd1 were used to build a sequence divergence tree. Consistent with many phylogenomics studies, the topology of the gene tree is similar but not identical to that of a species tree (Edwards et al., 2007). The presence of the Apcddc domains is indicated by the pink and green boxes. APCDD1 has two novel Apcddc domains (positions in human Apcdd1 protein: pink, amino acids 51-282; green, amino-acids 284-473) that resemble each other and may be an internal duplication. Some Apcdd1 proteins are shorter and have only one domain (Hemichordate phylum).

## Discussion

Here we report that Apcdd1 is a potent inhibitor of BMP signaling. We show that Apcdd1 blocks BMP signaling by binding to the type I BMP receptor, BmprIa, inhibiting the nuclear localization of pSmad1, the primary BMP effector. Overexpression of Apcdd1 represses the expression of BMP targets, and in *Xenopus*, chicken, and zebrafish embryos, interferes with major developmental processes including gastrulation, body axis formation, neural specification, and axon pathfinding. We previously showed that Apcdd1 also blocks canonical Wnt signaling by binding Wnt ligands as well as the LRP5 coreceptor. Thus, Apcdd1 provides a nexus for co-inhibition of both Bmp and Wnt signaling pathways. Protein expression and mathematical modeling of Apcdd1 inhibition of BMP and Wnt in hair follicles are consistent with the observed Wnt/BMP periodic sequential activation in mouse hair cycle (Lim and Nusse; Plikus et al., 2011) and with the complementary agent-based computational modeling that accompanied those experiments (Kandyba et al., 2013; Plikus et al., 2011; Plikus et al., 2008). Thus, Apcdd1 controls the duration of the phases of the hair follicle cycle by coordinating the timing of Wnt and BMP signaling activity (Fig. 5). These findings have a number of functional and evolutionary implications, as discussed below.

Only a few dual pathway inhibitors are known, and one (Sizzled) does not appear to be a *bona fide* dual Wnt/BMP inhibitor. While the ability to block both BMP and Wnt is a relatively novel feature of Apcdd1, other dual function inhibitors have been described, such as Cerberus, DAN, Gremlin, Coco,Wise/USAG-1 and Sclerostin, which are members of the CAN family of secreted peptides (Cruciat and Niehrs, 2013; Mulloy and Rider, 2015) that block BMP signaling via direct ligand-binding. In *Xenopus*, Cerberus also binds Wnt ligand, although this function is not conserved in mammals or birds. Of the CAN family, only Wise/USAG-1 and Sclerostin retain the ability to block both Wnt and BMP. Also, of note are members of the secreted Frizzled (sFRP) family of secreted peptides, which are related to the Wnt receptor Frizzled. Most sFRPs block Wnt signaling by direct ligand-binding (Cruciat and Niehrs, 2013). However, Sizzled represents a divergent sFRP member that blocks Bmp signaling. Sizzled, initially described as a Wnt antagonist (Salic et al., 1997), does not block Wnt but instead blocks BMP by stabilizing the Bmp antagonist Chordin (Bijakowski et al., 2012; Bradley et al., 2000; Bu et al., 2017; Lee et al., 2006; Muraoka et al., 2006; Yabe et al., 2003). Although Sizzled can no longer be considered a dual function inhibitor, its evolutionary derivation from sFRPs reflects an ancient relationship between BMP and Wnt regulatory mechanisms.

In contrast to the CAN and sFRP families, Apcdd1 stands alone as a new gene: unlike other inhibitors, Apcdd1 is not related to BMP or Wnt receptors or to any other gene, and has a highly conserved dual function as an inhibitor of both Wnt and BMP signaling. *Apcdd1* is a taxon-restricted gene present in a subset of Metazoan phyla, and contains a novel protein domain not found in other proteins, which leaves open the possibility that this gene arose not by copying other genes, but *de novo* or from long non-coding RNA (Carvunis et al., 2012; Tautz and Domazet-Loso, 2011). A similar example of a new gene arising by non-copying mechanisms at the origin of Metazoans and having important functions is the novel protein-coding gene Percc1 (Oz-Levi et al., 2019). *Apcdd1* appears to have evolved at roughly the same time as the BMP and Wnt signaling pathways, near the beginning of Metazoan evolution. Subsequently *Apcdd1* has been lost in a number of metazoan phyla. However, it is noteworthy that the Apcdd1 has been maintained in phyla in which both BMP and Wnt play key roles in establishing primary embryonic axes, whereas phyla do not utilize BMP for axial specification have lost Apcdd1 (Fig. 5A, Table S2). While this correlation warrants further investigation, our analysis of *Apcdd1* sequence and function illuminates how a taxon-restricted gene has been co-opted to coordinate the functional output of two major intercellular signaling pathways.

## Materials and Methods

### Xenopus laevis microarrays

4-cell stage embryos were injected marginally in each of the dorsal blastomeres with 10ng anti-*apcdd1* MO or control MO. *Apcdd1*-depleted embryos were also labeled on the ventral side with Nile Blue crystals, because some of the dorsally-depleted embryos displayed a delay in dorsal lip formation. At stage 10.5, when the dorsal lip is clearly marked, the vitelline membrane was removed with tweezers and the dorsal half sectioned with a hair knife(Vonica et al., 2011). 10 embryos were prepared for each condition (control and depletion) in three different experiments. RNA from each batch was purified by Trizol (Ambion) extraction and used for cRNA synthesis with the Gen Chip 3’IVT Express Kit (Affymetrix). Microarray (*Xenopus laevis* 2.0 Affymetrix Array, <u>GPL10756</u>) hybridization and data collection was performed by the Rockefeller University Genomics Center, and the data was analyzed using GeneSpring software. Complete microarray data was deposited in GEO.

### Xenopus gene expression studies

*Xenopus Apcdd1* cDNA was injected at the one-cell stage at a concentration of 150 ng/μl.

To evaluate the effect of *Apcdd1* on protein levels of TGFβ receptors, *Xenopus* embryos were injected in the animal poles of all blastomeres at-4 cell stage with RNAs for *alk3-myc* (100pg), *bmprII-HA* (200 pg), *alk4-HA* (250 pg), and *LacZ* (1 ng) as loading control, with or without *apcdd1-FLAG* (2 ng). Animal caps were collected at 4 cell stage in 1% NP40 buffer, deglycosylated, and run on NuPage 4-12% gels. Blots were incubated with the same antibodies as above, and with anti-beta-Gal chicken antibodies (1:5000), and visualized with ECL.

To evaluate the effect of *Apcdd1* on endogenous C-terminal phosphorylation of Smad1, 1ng *apcdd1* RNA was injected in the animal pole of all blastomeres at the 4-cell stage. Embryos injected with *chordin* RNA (1 ng) served as controls for the inhibition of BMP-dependent Smad1 phosphorylation. Animal caps were dissected at stage 11 and lysed in the presence of protein phosphatase inhibitor cocktails (Sigma). Western blots were stained with rabbit anti-Smad1 antibody (1:1000, Cell Signaling) and rabbit anti-pSmad1 (C-terminal) antibody (1:2000, from E. Laufer, Columbia U.).

#### Expression plasmids

p64T *Xenopus* CA-ALK3 (A. Suzuki, Hiroshima U.), pCS2+ *Xenopus* ALK4-HA (A.V), pCS2-Xnr1 (C. Wright, Vanderbilt U.), pCS2 BMP7 (A. Brivanlou, Rockefeller U.), pSP64T BMP4 and p64T BMP2 (G. Thomsen, SUNY Stony Brook), pCS2 (*Xenopus*) BMPRII-HA (M. Asashima, Tokyo U.), pcDNA3 (human) ALK3-Myc (E. Laufer, Columbia U. Coll. Phys. & Surg.), pSP35T Chordin (E. Amaya, U. Manchester), pGEM5Zf(-) Wnt8 (R. Harland, U.C. Berkeley), pCS2 *apcdd1* (*Xenopus*) (A.V.), pBS *APCDD1, APCDD1 FLAG*, *APCDD1 L9R*, and pCXN2.1 *APCDD1 FLAG* (Y. Shimomura, Niigata U.).

### Embryo manipulation, RNA *in vitro* transcription, and RNA *in situ* hybridization

*Xenopus* embryos were obtained by *in vitro* fertilization and cultured in standard solution (MMR). RNAs and MOs were injected on the dorsal side (bilaterally) or animal side (both ventral blastomeres for BMP transcription, all 4 cells for phospho Smad1 Western blot). For rescue of anterior structures, *apcdd1* MO (20ng) were injected alone or together with 1 ng *Bmp2*, *Bmp4*, *Bmp7*, *admp* MO in the marginal zone of both dorsal blstomeres at the 4-cell stage and collected at stage 34. At the specified stages, according to Nieuwkoop and Faber, embryos were fixed for *in situ* hybridization, or extracts prepared for transcription or protein gels. RNA for injection and *in situ* hybridization was prepared with mMessage Machine with SP6, T7 or T3 RNA polymerases (Ambion), and T3 or T7 polymerase (Roche), for *in situ* probes with digoxigenin or FITC labeling mix (Roche), respectively. Morpholino oligonucleotides against *Xenopus* BMPs were: *admp* MO, 5’ – GGTCCATCTCATCAAGCTGCAGCTC-3’(Reversade and De Robertis, 2005), Bmp2 MO, 5′-GATCCCAGCGACC ATTGTCAACCTG-3′; Bmp4 MO, 5′-CAGCATTCGGTTACCAGGAATCATG-3′; Bmp7 MO, 5′-TTACTGTCAAAGCATTCATTTTGTC-3′(Reversade et al., 2005), and were co-injected with Apcdd1 MO at 1 ng /dorsal blastomere.

### Transcription assays in embryos and cells

*Xenopus* embryos: transcription assays for the Wnt pathway were done as in (Shimomura et al., 2010), using the TOP-FLASH reporter. For BMP-dependent transcription in Apcdd1-depleted embryos, MO against Apcdd1 were injected in the animal poles at 2-cell stage (20 ng in each of 4 injections), followed by injection of *BRE* reporter (50 pg) and *Bmp2* RNA (50 pg) in the animal poles of ventral blastomeres at 4-cell stage. For inhibition of Bmp activity by overexpressed *Apcdd1*, *BRE* reporter and *Bmp2* RNA were coinjected with RNA of wild-type *Apcdd1* or L9R mutant (Shimomura et al., 2010) *Apcdd1* (1.5ng) in the animal poles of both ventral blastomeres at 4-cell stage. For constitutive Alk3 transcription, embryos were collected at stage 10.5, four embryos/assay, in triplicate, and each experiment was repeated a minimum of 3 times. Figures show a typical transcription assay for each experiment.

Cells were cultured in 24-well dishes, transfected with DNA (200 ng *Bmp4* expression vector, 100 ng BRE reporter, 10 ng TK-Renilla control plasmid, 50 or 150 ng wild-type and L9R mutant *APCDD1*), and collected after 36 h. Luciferase activity was measured with the Dual Luciferase kit (Promega), in triplicates, and the experiment was repeated three times. The figure shows a typical experiment.

Luciferase transcription assays, in *Xenopus* and NIH 3T3 cells, for the Wnt pathway were carried out as in (Shimomura et al., 2010), using the TOP-FLASH reporter, while transcription assays for the BMP pathway used the *BRE* reporter activated by Bmp2 (Hata et al., 2000).

### Cell culture immunoprecipitation analysis

CHO cells and NIH 3T3 cells were grown in standard growth medium and transfected with Fugene (Roche) or Lipofectamine 2000 (Life Sciences).

Plasmids expressing human Alk3 Myc (500ng), Bmrp2 HA (1µg), and human Apcdd1 FLAG (500pg) were transfected in CHO cells grown in 6-well plates. Cells were collected after 36 hours, lysed in 1% NP-40 buffer with added 1% Triton, 10% glycerol, 1mM EDTA, and protease inhibitors (Roche), and immunoprecipitated with either anti-FLAG (M2, Sigma), or anti-Myc beads (Millipore). After washes, immunoprecipitated receptors were deglycosylated with PNGase F (New England Biolabs), before resuspending in LDS sample buffer and running on NuPAGE 4-12% Bis Tris precast gels, using MOPS buffer (Invitrogen). Vector-transfected cells served as control. Proteins were transferred to PVDF membranes, incubated with mouse anti-HA (1:1000, Covance), rabbit anti-Myc (1:1000, Millipore), and rabbit anti-mouse Apcdd1 (1:5000, from Dritan Agalliu, UC Irvine), then with HRP-labeled secondary antibodies before staining with ECL (Amersham).

### Immunofluorescence for BMP and Wnt signaling effectors in the presence of *Apcdd1*

NIH 3T3 were cultured in 8 well chamber slides with cover (LabTek II) transfected with empty vector or *Apcdd1-eGFP*. Cells were starved overnight in Optimem (Gibco) before addition of growth medium with or without recombinant signaling molecules (20 ng/ml human BMP2, 100 ng/ml Wnt3a, separately and together, R&D) for 2 h before fixation. Primary antibodies were mouse monoclonal anti-β-catenin (clone 5H10) and anti-Phospho-Smad1 (C-terminal) as above, and secondary antibodies were anti-mouse Alexa 647 and anti-rabbit Alexa 594 anti-rabbit antibodies (ThermoFisher).

The images were acquired with a confocal Zeiss LSM 510 confocal microscope. Images were acquired sequentially for each channel such that no image region was saturated or below detection The images were pseudocolored in Image J as follows: Apcdd1 in red, pSmad1 or β-catenin in green, DAPI (to visualize the nuclei) in blue. Channels were split in Image J and the level of pSmad1, β-Catenin, and DAPI were measured in the nucleus, whose contours were determined using DAPI. GFP levels were measured in the whole cell, using the β-catenin peri-membrane signal to determine the outline of the cell. For each of the eight groups of cells (two groups: transfected or not-transfected cells; for each group, there are 4 subgroups: no ligand, BMP2, Wnt3A, BMP2 + Wnt3A), at least 40 cells were imaged. For every cell, the background was determined by using the same mask to measure the signal in an empty area of the slide. The adjusted signal was reported as *S_adjusted_ = (S_cell_ -S_background_)/S_background_*. Within the figures (Fig. 1A-B), individual channels were merged and displayed as follows: DAPI (blue), fluorescent protein reporter of Apcdd1 (red), pSmad1 (green, Fig. 1A), β-catenin (green, Fig. 1B). The images were pseudocolored in Image J as follows: APCDD1 in red, pSmad1 or β-catenin in green, DAPI (to visualize the nuclei) in blue.

### Immunofluorescence for the effect of inducible expression of Apcdd1 on Alk3 expression in NIH 3T3 cells

The T-Rex system (Invitrogen) was used for tetracycline-dependent expression of human *Apcdd1*. NIH3T3 cells cultured in 8 well slides were transfected with 50 ng *ALK3-Myc*, 50 ng pcDNA 4/TO *apcdd1*, and 100 ng 6/TR (expressing the TET repressor) plasmids, with empty vectors, *ALK3-Myc* plasmid alone, and no-tetracycline wells as controls. Slides were fixed at time 0, and 1, 2, 4, 8, 12 h after tetracycline addition, stained with mouse anti-Myc (1:200) and rabbit anti-Apcdd1, secondary anti-mouse Alexa 488, anti-rabbit Alexa 594 (Life Technologies) and subjected to confocal microscopy.

### Morpholino injections

To knockdown *apcdd1* in zebrafish, embryos were co-injected with 10 ng/nl *apcdd1* translation blocking morpholino (5’GAATAAGTCTCTCCCCAGCCATGAT3’) along with 5 ng/nl *p53*-MO, in a total volume of 1 nl/embryo. *p53*-MO was included to block non-specific cell death, though cell death was negligible even without *p53*-MO.

### Pharmacological inhibition of Bmp

Dorsomorphin (Sigma, P5499) stock was dissolved in DMSO at a concentration of 10 mM. It was subsequently diluted to a concentration of 75 μM in embryo medium and added to zebrafish embryos at 10 hpf until fixation.

### Apcdd1 probe synthesis

After extraction of total RNA, cDNA was synthesized using oligo dT primer using Invitrogen kit. Zebrafish *apcdd1l* cDNA was PCR amplified and cloned into pCS2+. The probe was subsequently synthesized using sense strand as a template.

### Chicken immunohistochemistry

Antibody staining and RNA *in situ* hybridization histochemistry were performed on 15 μm transverse sections from Hamburger and Hamilton (HH) stage 14-24 (Hamburger and Hamilton, 1992) chicken spinal cords as previously described (Augsburger et al., 1999). Fluorescence images were collected on a Carl Zeiss LSM510 confocal and Axioplan 2 microscope. Images were processed using Adobe Photoshop CS4.

An RNA probe for *in situ* hybridization against the 3’ UTR of the chicken *APCDD1* gene was prepared using the following primers: forward, 5’-GAG <u>ATT AAC CCT CAC TAA AGG GA</u>T GCT GCC TCA AAA ACA GAT G -3’, reverse 5’-CAG CCT TGA GGC CTT TAC TG -3’. The underlined region denotes a T3 polymerase site embedded in the primer sequence. The target sequence was amplified from HH stage 16/20/24 chicken spinal cord cDNA by PCR. Qiaquick (Qiagen) purified products were used in an *in vitro* transcription reaction using the Roche DIG RNA labeling kit.

Antibodies were used to detect several proteins: <u>rabbit</u> anti Lhx2/9 (pan Lh2a/b), 1:1000 (Liem et al., 1997); rabbit Islet1/2 (K5), 1:2000 (Tsuchida et al., 1994), rabbit anti-Pax2, 1:250 (Invitrogen); <u>mouse</u> anti-GFP at 1:1000 (3E6, Invitrogen). Species-appropriate Cyanine-3 and Fluorescein conjugated secondary antibodies were used (Jackson ImmunoResearch Laboratories).

### Expression constructs and *in ovo* electroporation

Fertile White Leghorn eggs (McIntyre Poultry) were incubated to Hamilton and Hamburger (HH) stages 14/15 to examine cell fate specification phenotypes or stages 16/17 to axon outgrowth phenotypes. The following expression constructs were electroporated *in ovo* into the developing neural tube as described (Briscoe et al., 2000): CMV::*gfp* (pCIG, 0.5μg/μl), Math1::farnesylated (f) *gfp* (0.2μg/μl-0.5μg/μl), CMV::*APCDD1* (0.2μg/μl) and the resulting eggs incubated until HH stages 22/23. Cell fate and dI1 axon phenotypes were quantified as described (Hazen et al., 2011; Phan et al., 2010), respectively. All statistical analyses were performed using a two-tailed Student’s *t*-test.

### Statistical analysis

Using Excel, several parameters of the distributions of the adjusted fluorescence signals were computed for each condition (mean, median, mode, standard deviation, coefficient of variation, standard error of the mean, skew of the distribution, kurtosis). Since the distributions were essentially normal, the different conditions were compared using Student’s two-tailed t-test, adjusting the p-values for multiple comparisons using the Bonferroni correction.

### Hair follicle immunochemistry and immunofluorescence

Swiss-Webster mice were selected to have a clear view of the hair matrix without melanin pigments. Mice were sacrificed at P31, during mid-anagen, and hair follicles examined were in Anagen V-VI, according to previous criteria (Muller-Rover et al., 2001). Dorsal skins of the mice were prepared for frozen sections and paraffin-embedded sections. Frozen sections 8 µm thick were post-fixed with 4% paraformaldehyde, blocked in 2% fish-skin gelatin, and incubated with primary antibodies to APCDD1 (1:1000, gift of Dr. Dritan Agalliu), pSMAD-1/5 (1:250, gift of Drs. Thomas Jessell and Ed Laufer), P-cadherin (1:500, Invitrogen 13-2000Z) and Ki67 (1:250, Santa Cruz sc-7846) at 4°C overnight, and counterstained with secondary antibodies. Paraffin-embedded sections were rehydrated in graded ethanol, subject to sodium citrate antigen retrieval, peroxidase blocking with 3% hydrogen peroxide, and incubated with primary antibody to β-catenin (1:100, BD Transduction 610153) at 4°C overnight, followed by a HRP-conjugated anti-mouse secondary antibody (Vector Labs, MP-7500), and developed with ImmPACT NovaRED kit (Vector Labs, SK-4805) according to manufacturer’s instructions. Immunofluorescence results of frozen sections were viewed on a Carl Zeiss LSR Confocal Microscope, and all bright-field images of paraffin-stained sections were obtained a Zeiss Axioplan2 Microscope.

### Mathematical modeling of signaling pathway activation. Short description of the mathematical model

The model applies to the dynamics of BMP and Wnt signaling within hair follicles. In adult hair follicles, BMP and Wnt signaling are coordinated, are activated with the same period length, and are out of phase (Kandyba et al., 2013; Plikus et al.; Plikus et al., 2008). The coordination of BMP and Wnt activation dynamics in adult mouse hair follicles has previously been modeled with an agent-based model based on cellular states and not on the proteins involved in the process (Kandyba et al., 2013; Plikus et al.; Plikus et al., 2008). Our model of the same process in hair follicles takes a complementary approach, since it is a numerical model based on coupled differential equations used to evaluate protein level dynamics. “It assumes Michaelis-Menten kinetics, following a framework that is commonly used in the context of signaling and metabolic pathways (see, for example, Sections 1.5 and 7.2 of (Keener and Sneyd, 2009), and (Suel et al., 2007)).”

Both BMP and Wnt are treated as model inputs, denoted by f and g (respectively) in the model equations, and measured in arbitrary units. We consider four cases according to whether the inputs are constant or time-dependent, periodic functions:

I. The two inputs BMP and Wnt are both constant and continuous.
II. The two inputs BMP and Wnt are both periodic (either in-phase or out-of-phase by 1/4 of the period).
III. BMP is constant, Wnt is periodic.
IV. BMP is periodic, Wnt is constant.

In Fig. 4D, we show configuration **II**, in which BMP and Wnt ligand inputs are both periodic and out-of-phase by 1/4 of the period.

We make the following notations: y_1_=APCDD1, y_2_=BmpR-total, y_3_=BmpR-activated, y_4_=pSmad1, y_5_=WntR-activated, y_6_=β-catenin, f=Bmp, g=Wnt (f and g are inputs), c=WntR-total. We are considering the possibility that the BMP pathway may inhibit the Wnt pathway.

We plot pSmad1 (in blue) and β-catenin (in red) against time. In Fig. 4D, the plots correspond to the following 5 cases:

**Case i**, no inhibition coming from APCDD1, as in null mutants;
**Case ii**, low inhibition coming from APCDD1, as in mutant APCDD1;
**Case iii**, full inhibition coming from APCDD1, as in wild type APCDD1;
**Case iv**, full inhibition coming from APCDD1, as in wild type APCDD1; additionally, there is repression of the Wnt pathway by the BMP pathway.
**Case v,** full inhibition coming from APCDD1, as in wild type APCDD1; additionally, there is repression of the Wnt pathway by the BMP pathway; in this case, APCDD1 is also induced by Wnt signaling.

Here are the parameters that we use:

- η= Gaussian noise with standard deviation 2 ×10^-4^ and η’= Gaussian noise with standard deviation 2×10^-5^,
- basic levels *c*=0.035, *c_1_*=0.016, *c_2_*=0.0054, *c_3_*=0.0001, *c_4_*=0.004,
- parameters *a_1_, a_2_, a_3_* are coefficients for various activating terms as follows: *a_1_=6, a_2_=5, a_3_=4,*
- parameters *b* and *b_1_* are coefficients for various inhibiting terms as follows: *b*=0.1 and *b_1_*=0.0003 for low inhibition (**case ii**), *b_1_*=0.0008 for full inhibition (**cases iii, iv,v**).

For Figure 4D, the equations for the **Case i** are:

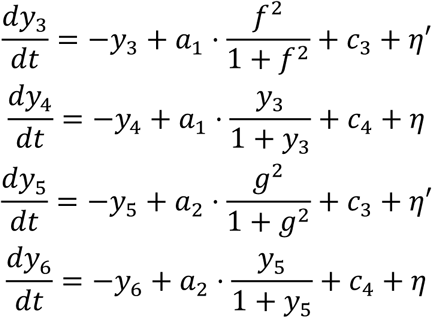

The equations for the Figure 4D **Cases ii, iii** are:

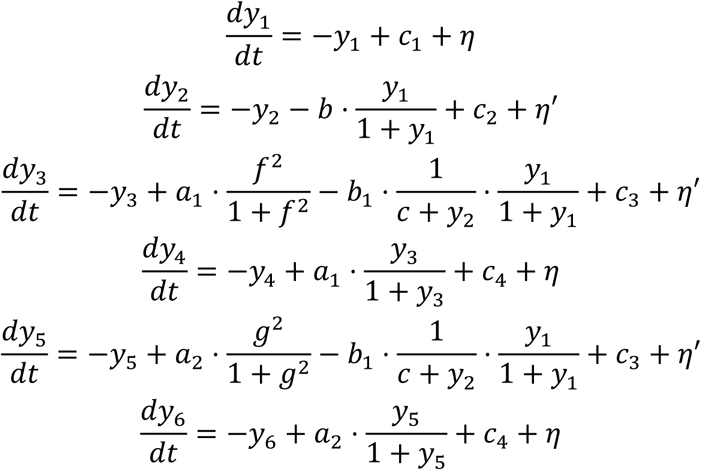

The equations for the Figure 4D **Case iv** are the same as in the previous case, except for the last line, which reads:

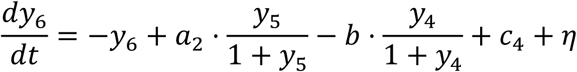

The equations for the Figure 4D **Case v** are the same as in the previous case, except for the first line, which reads:

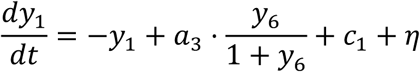

We note that we studied many more formal cases which are not shown here.

In Fig. S6, similarly, we plot pSmad1 (in blue) and β-Catenin (in red) against time. The plots correspond to the following 2 cases:

In Fig. S6, **Case A** (left column in Fig. S6A) b_1_=b_2_=b=0, no inhibition coming from APCDD1, with mutual activation between the pathways;

In Fig. S6, **Case B** (right column in Fig. S6A) APCDD1 is produced by the Wnt branch and also added exogenously, full inhibition from APCDD1, with mutual activation between the pathways.

We note that we studied many more formal cases which are not shown here.

Here are the parameters we use:

- basic level a_0_=10^-3^ and η= Gaussian noise with standard deviation 2·10^-4^
- basic level a’_0_=10^-4^ and η’= Gaussian noise with standard deviation 2 ·10^-5^
- basic level a’’_0_=8·10^-4^ and η’’= Gaussian noise with standard deviation 10^-4^
- parameters K and a_1_ up to a_6_ are coefficients for various activating terms as follows: a_1_=6, a_21_=4, a_25_=7, a_26_=1, a_3_=0.015, a_4_=0.0053, a_5_=0.6, a_6_=0.2, b=0.1, K=0.035
- the parameters b_1_ and b_2_ are coefficients for inhibition terms (b_1_, b_2_) = (0.0006, 0.0003).

The equations for In Fig. S6, **Case A** are:

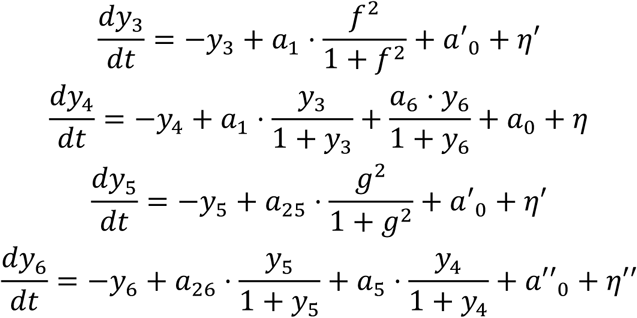

The equations for In Fig. S6, **Case B** are:

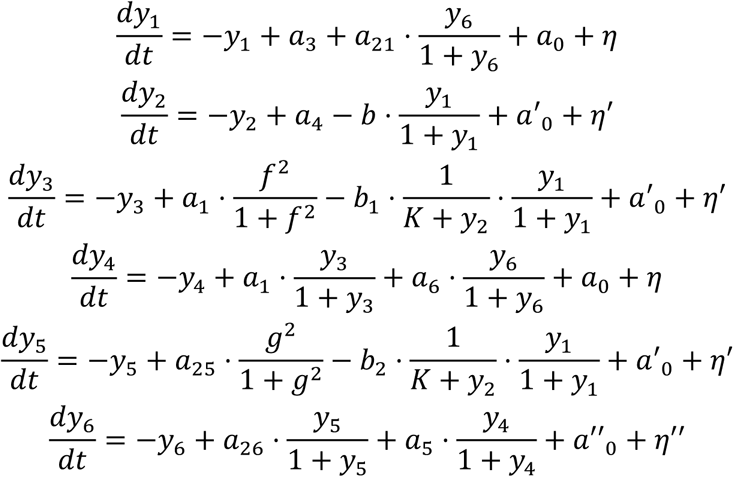

### Apcdd1 synteny, detection of conserved domains, and gene divergence analyses

For synteny analyses, within genomes with Apcdd1 annotated, we looked at the nearest three upstream gene neighbors and three downstream gene neighbors using the UCSC Genome Browser and NCBI. To determine whether conserved domains exist within Apcdd1 proteins, we performed BLASTP queries (Altschul et al., 1990) using human APCDD1 with an E-value of 10^-20^. During alignment, we noted the presence of a novel domain, which was further analyzed with HHrepID (Biegert and Soding, 2008). Consensus Apcdd1 domain sequences for mammals were obtained by aligning the mammalian sequences with CLUSTALW in MEGA 5 (Tamura et al., 2011) followed by WebLogo 3 analysis (Crooks et al., 2004) to obtain the consensus sequence. In some proteins, the domain was found to be duplicated. Further BLASTP searches with the domain 1 and domain 2 mammalian consensus sequences retrieved six bacterial sequences, including *Stigmatella, Methanobacterium*, and *Spirosoma*. We note that this domain has also been recently annotated by the Pfam (El-Gebali et al., 2019) database curators, who named it Apcddc. The Pfam Apcddc domain boundaries correspond to the boundaries we had previously identified. Moreover, using the Apcddc domain to query BLASTP and HMMER (Eddy, 1998) retrieves only Apcdd1 and *Apcdd1*-like genes, indicating this domain occurs only in *Apcdd1* genes. Furthermore, a phylogenetic tree of all collected sequences was constructed using MEGA 5.21 (Tamura et al., 2011) with alignment of the protein sequences with CLUSTALW using standard parameters (gap opening penalty 10, extension 0.1). A Neighbor-Joining phylogenetic tree was constructed using the Maximum Composite Likelihood Model with pairwise deletion and bootstrapped 5000 times. The initial tree was used to distinguish *Apcdd1* sequences (closest to human APCDD1) from *Apcdd1*-like sequences.

To detect remote similarity, *Apcdd1* sequences were selected for further analysis and a subsequent alignment. A BLASTP query against the NCBI non-redundant database was used to retrieve *Apcdd1* protein sequences that are similar to human APCDD1 with an E-value of 10^-3^, allowing 200,000 as the maximal number of hits. After removing isoforms, 441 sequences remained. Low-quality alignments were removed, including a small number of bacterial sequences.

Sequence alignments of the APCDD1 sequences were performed in 30 chordate species for the protein sequences most similar to human APCDD1. MUSCLE (Edgar, 2004) was used with ClustalW sequence weighting and UPGMB clustering. PartitionFinder2 (Lanfear et al., 2017) was used to identify the optimal partition scheme and evolutionary models. Using these results, RAxML (Stamatakis, 2014) was used to generate a maximum likelihood gene tree using the PROTGAMMAWAG substitution model and 1000 bootstraps. FigTree was used to visualize the gene tree (Fig. S6B).

### Apcdd1 protein sequence and secondary structure analysis

To analyze the content of secondary structure elements within Apcdd1 proteins, we used bioinformatics algorithm collections PredictProtein (www.predictprotein.org) and HMMER (hmmer.org) (Eddy, 1998; Yachdav et al., 2014). Within 441 Apcdd1 proteins from different Metazoan species, we scored sequence features and predicted secondary structural properties according to over 25 criteria. These criteria include disorder, content of α-helices and β-strands, number of binding sites to other proteins or to DNA, subcellular localization, propensity to aggregate, propensity to form prions, presence of transmembrane domains, presence of signal peptides, presence of Pfam domains, amino acid composition.

## Acknowledgments

V.L. and A.M.C. acknowledge funding support from NIH (grants: P30AR044535 to David Bickers, R01AR052579 to A.M.C). V.L. thanks Marc W. Kirschner and acknowledges funding support from National Institutes of Health (NIH grants R01 HD073104 and R01 HD091846, to M.W.K.). V.L. and L.I. acknowledge partial funding support from the Principal’s Interdisciplinary Fund of the University of Aberdeen and the National Science Foundation (grant number PHY11-25915 to U.C.S.B., K.I.T.P.). Funding support was provided to A.H.O.L. by the NIH National Child Health and Development Institute (K12 HD052896) and by a Boston Children’s Hospital Career Development Fellowship. A.V. was supported by NIH grant R03 HD057334 and is grateful to Dr. Jean Gautier at the Institute of Cancer Genetics, Columbia University in New York City, for access to the lab *Xenopus* facility. Zebrafish experiments were supported by NIH-NIDCD grant R01-DC03806. S.J.B. and K.P. acknowledge funding support from NIH/NIHDS (R01: NS085097) and from the UCLA Broad Stem Cell Research Center. V.L. thanks Wenzhe Ma and Harry Burgess for insightful comments. V.L. and A.M.C. thank Yutaka Shimomura for his kindness and inspiring style.

## Author contributions

A.V., V.L., and A.M.C. conceived of the project. A.V. performed biochemistry, frog, and cell culture experiments. K.P. and S.J.B. performed the chicken experiments. N.B., J.G., and B.R. performed zebrafish experiments. V.L. performed cell imaging experiments. V.L. and L.I. did the mathematical modeling. E.W. and G.M.D. performed hair follicle immunohistochemistry. A.H.O.L., V.L., A.K., J.W.C. and J.A.W. performed phylogenetic and protein domain analyses, and statistical analyses. V.L., S.J.B., J.A.W., B.R., A.V., and A.M.C. wrote the paper. All authors read and edited the manuscript.

## Supplementary Data

**Data S1. Supplementary Table 1 with array genes** is presented separately as an Excel file.

**Supplementary Table 2.**
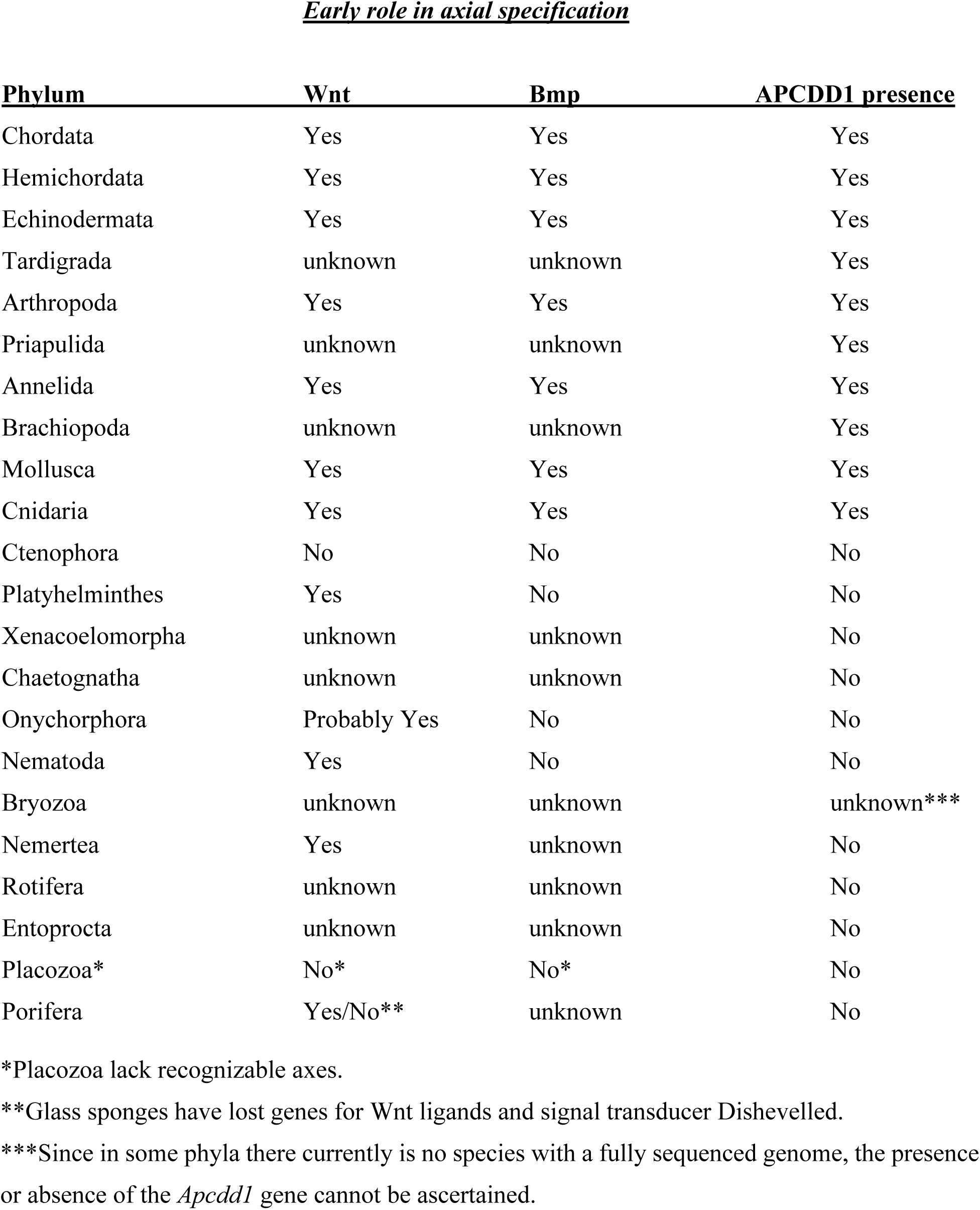
The presence of the *Apcdd1* gene correlates with the involvement of Bmp signaling in axial specification. The involvement of Wnt activity and Bmp activity is listed in Metazoan phyla.

**Figure S1.**
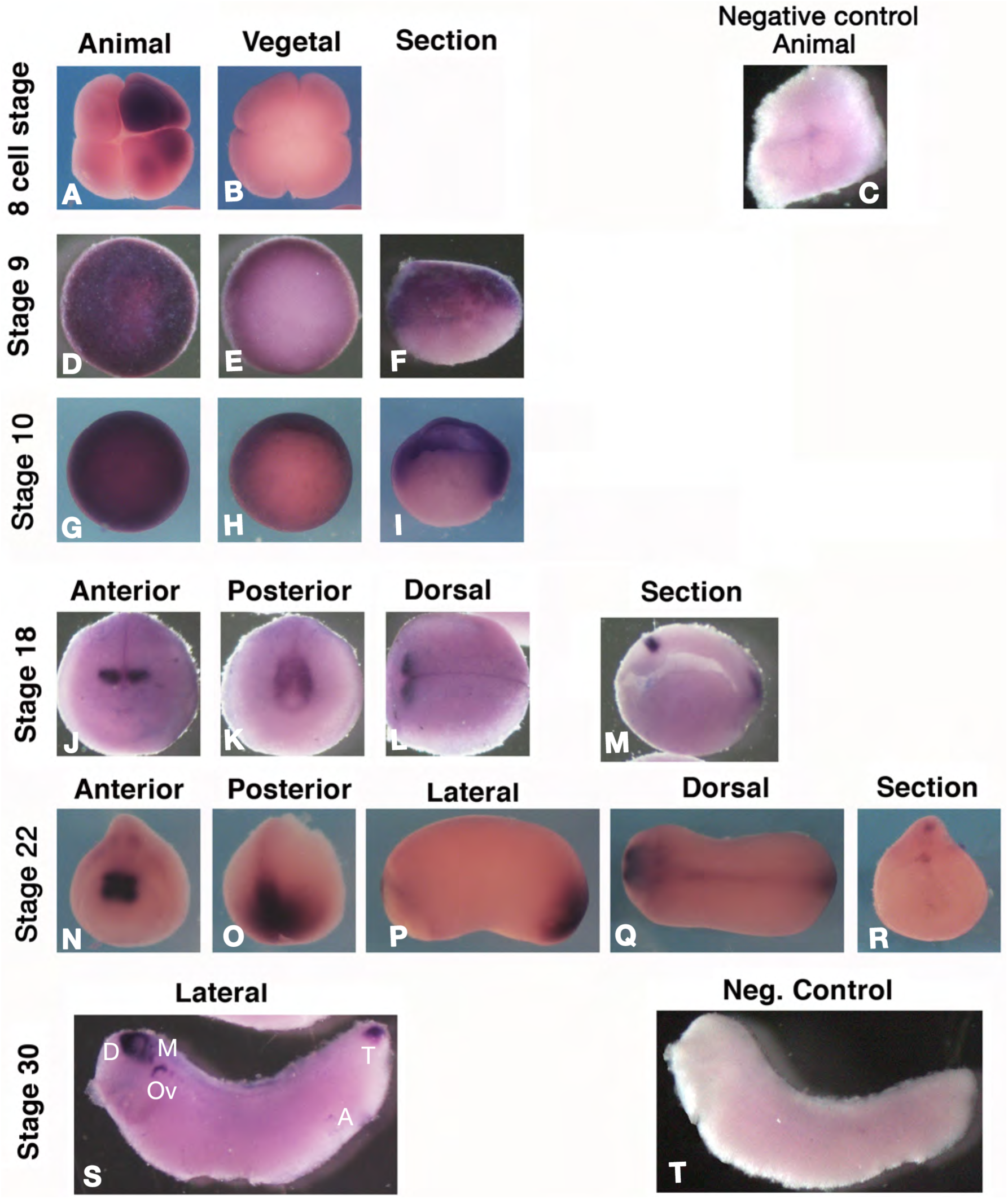
Expression of *Xenopus Apcdd1*. **A-T.** RNA *in situ* hybridization for *Xenopus Apcdd1*. Maternal expression at 8 cell stage (A, B) is localized in the animal pole and is frequently unevenly distributed. In late blastulas (stage 9, D-F), the expression is animal and marginal (arrows in F, blastocoel in outlined in F). At the start of gastrulation (stage 10, G-I) *Apcdd1* RNA is present in animal and marginal cells, including the dorsal lip with the Spemann organizer (arrow in H, I, blastocoel is outlined in I). In neurula embryos (stage 18, K-N) and tailbud (stage 22), expression is limited to diencephalon and midbrain precursors (J, K, M, N, Q), and the tailbud (K, M, O, P, Q). Arrowhead in the transversal section (R) indicates ependymal precursors. In tadpoles (stage 30, S) expression closely mirrors that in mouse embryos. D – diencephalon; M – midbrain; Ov – otic vesicle; T – tailbud; A – anus. Bracket in S indicates the branchial arches. For control *in situ* hybridization (C: 8-cell stage; T: tadpole), the RNA probe used was *LacZ*. The scale bar in A is 0.3 mm (all embryos are shown at the same scale)

**Figure S2.**
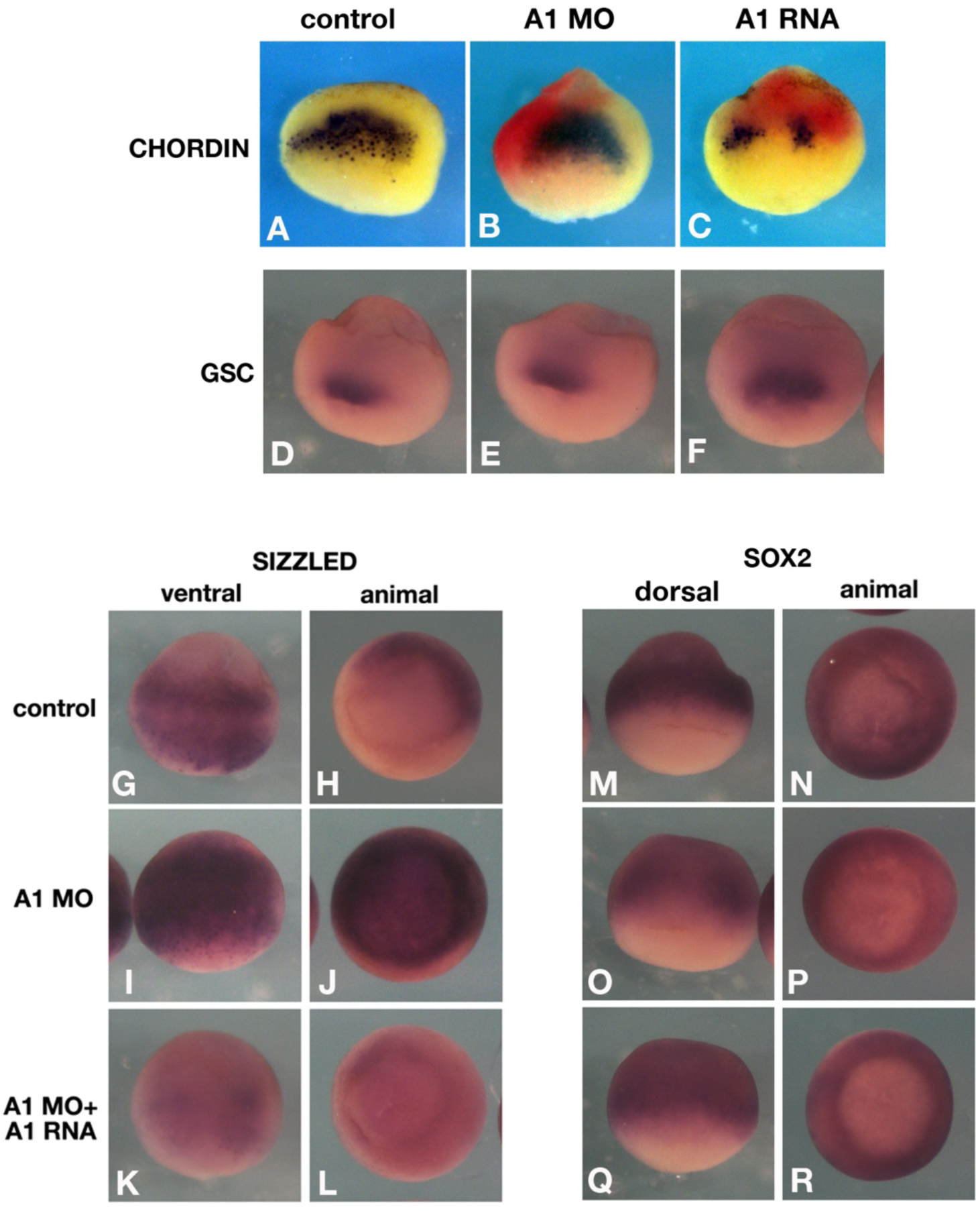
*Apcdd1* regulates expression of both positive and negative targets of BMP signaling in *Xenopus* embryos. *In situ* hybridization of morpholino-depleted (A1MO), or Apcdd1 overexpressing embryos. Embryos were injected at the 4-cell stage dorsally (A-C, double *in situ* hybridization for *chordin* and *LacZ* RNA as injection marker, in red), or 1 cell stage (D-R). Control embryos were injected solely with *LacZ* RNA (1 ng). **A-F.** Spemann organizer genes: *chordin* is not affected by Apcdd1 depletion (B) (n=18), but inhibited by *Apcdd1* RNA (A1 RNA) overexpression (n=23), consistent with the inhibition of the maternal, dorsal Wnt pathway; *gsc* is unaffected by either treatment (n= 15 and 12, respectively, E, F). **G-L.** The ventral marker and BMP target *sizzled* is expanded by Apcdd1 depletion (n=23, I, J), and restored by expression of a MO-resistant *Apcdd1* RNA (n=21, K, L). **M-R.** The early neural marker and negative BMP target *sox2* is inhibited by Apcdd1 depletion (n=27, O, P, arrows in O indicate a hole in the ventral expression pattern), and partially restored by expression of *Apcdd1* RNA (n=17, Q, R).

**Figure S3.**
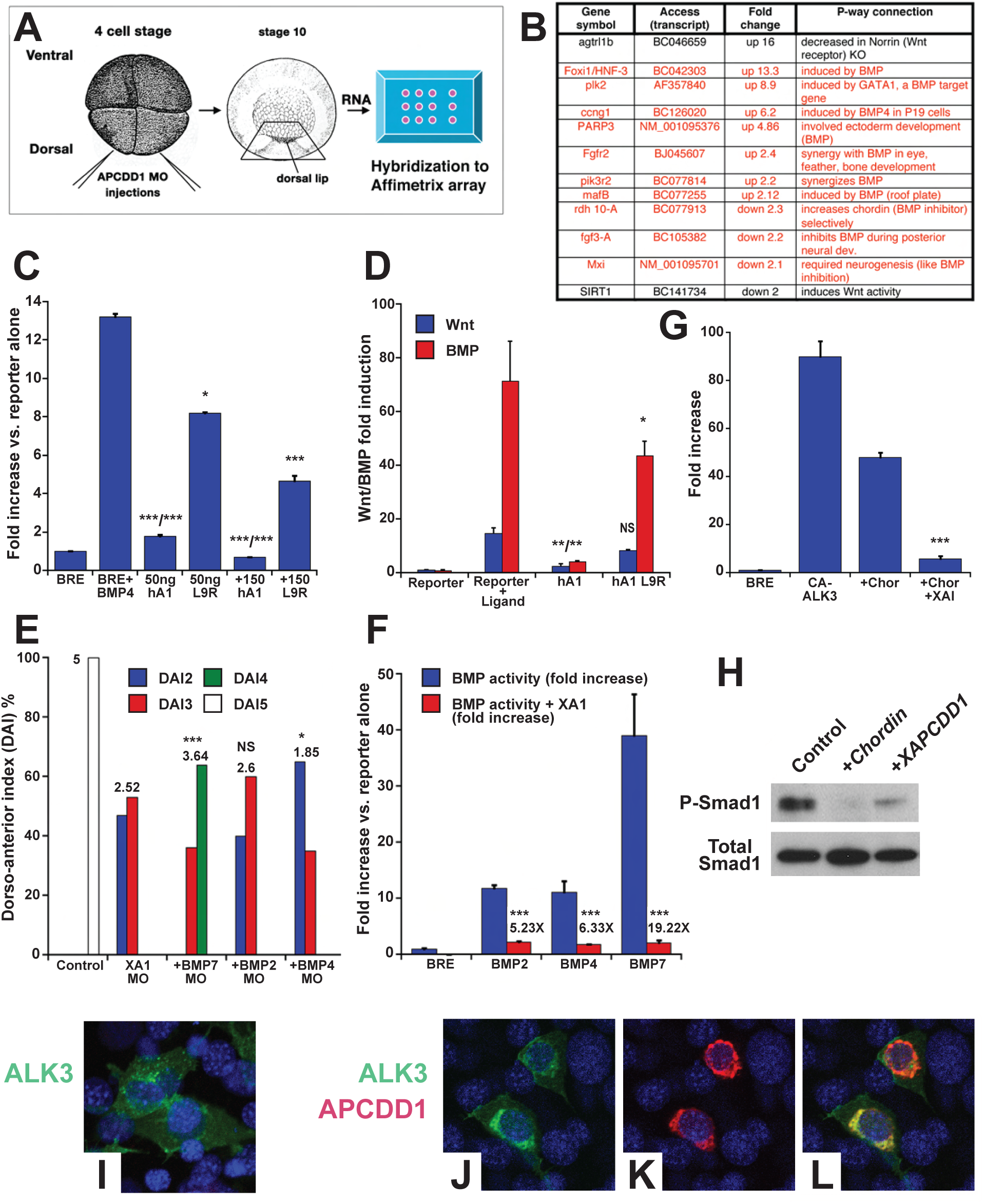
Apcdd1 inhibits BMP signaling in embryos and in cultured cells. **A.** Microarray detection of genes dependent on *Xenopus laevis Apcdd1* (XA1) in *Xenopus* embryos. Apcdd1 protein was depleted with morpholino oligonucleotides in the dorsal part of the embryo injected at the 4-cell stage, and dorsal tissue containing the Spemann organizer was recovered at stage 10. Extracted RNA was used to identify changes in gene expression versus control embryos in Affymetrix microarrays. **B.** Protein coding genes with more than 2-fold variation in XA1-depleted embryos. Genes shown were verified by quantitative PCR. Red: BMP-regulated genes; black: Wnt-regulated genes. **C.** Effect of human *APCDD1* (hA1) on BMP activity induced by recombinant BMP4 in NIH3T3 cells. The L9R mutant form of APCDD1 is significantly less active. **D.** Apcdd1 inhibits both the Wnt and BMP pathways induced by exogenous *Wnt8* and *Bmp2* RNA in *Xenopus*. The APCDD1-L9R mutant has a considerably reduced effect on both pathways. **E.** Quantification of XA1 depletion and rescue experiment (Fig. 1 E-I) results in a percentage of embryos with a specific dorso-anterior index (DAI) (Kao and Elinson, 1988). Numbers on top indicate the average DAI for the batch of embryos. **F.** Induction of BMP signaling by different BMP ligands is differentially suppressed by XA1. **G.** The combined action of chordin and *Xenopus* Apcdd1 is sufficient to suppress BMP-dependent reporter gene activation by constitutively active form of the type I BMP receptor (caBmprIa). **H.** *Xenopus* Apcdd1 is sufficient to reduce Smad1 C-terminal phosphorylation induced by endogenous BMPs in *Xenopus* animal caps. Chordin is a positive control for BMP inhibition. **I-L.** Co-expression of type I BMP receptor BMPRIA and tetracyclin-inducible APCDD1 expression in CHO cells causes displacement of BMPRIA after 12 h. Statistical significance: * p < 0.05; ** p < 0.01; *** p < 0.001, Student’s t test.

**Figure S4.**
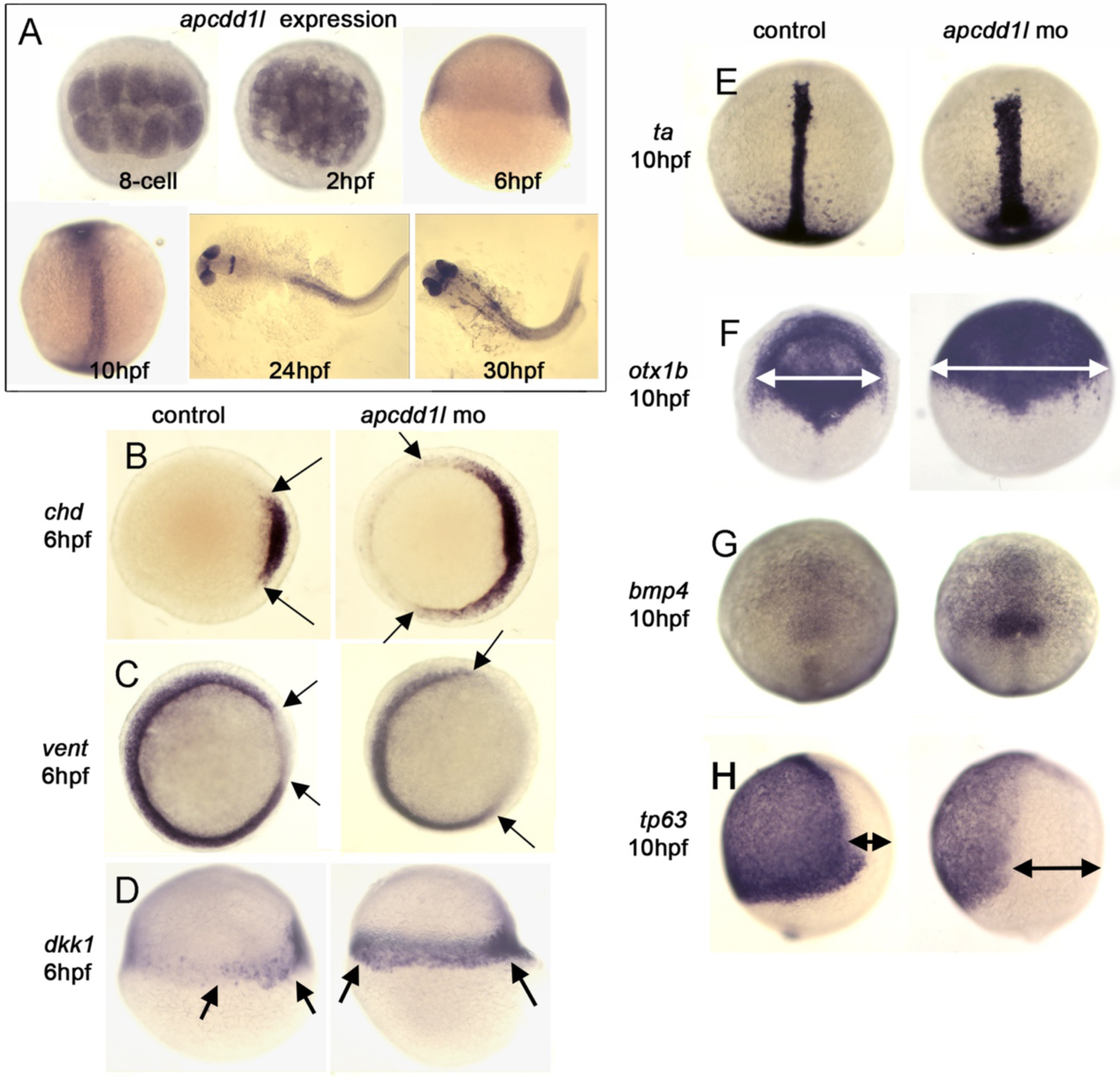
Expression of zebrafish *apcdd1* and effects of functional knockdown on embryonic patterning. **A.** The presence of maternal *apcdd1l* mRNA, confirmed by RT-PCR (not shown), is seen in all cells in the early blastula (animal pole views). By early gastrulation (6 hpf), *apcdd1l* is only weakly expressed with preferential staining in dorsal tissues (lateral view, dorsal to the right). By the end of gastrulation (10 hpf), *apcdd1l* is expressed throughout to the dorsal axis (dorsal view and lateral view, anterior up). At 24 and 30 hpf, *apcdd1l* is expressed in the eyes, restricted regions in the brain, and dorsal tissues in the trunk and tail. **B-H.** Domains of expression of various markers of gastrula stage embryos. Expression of *chd* in the organizer and *vent* in ventrolateral mesoderm (B, C, animal pole view) and *dkk1* in the organizer (D, lateral view). The organizer is expanded laterally at 6 hpf in *apcdd1l* morphants, and there is a corresponding contraction of ventrolateral tissue. At 10 hpf, expression of *brachyury-a* (*ta*) in the notochord (E) and *otx1* in the anterior neural plate (F) (dorsal views, anterior up) are expanded in *apcdd1l* morphants, whereas expression of ventral marker *tp63* in epidermal ectoderm (H, lateral view) is contracted. At this time, expression of *bmp4* at the ventral midline (G, dorsal view) begins to upregulate and expand in *apcdd1l* morphants, presaging subsequent ventralization.

**Figure S5.**
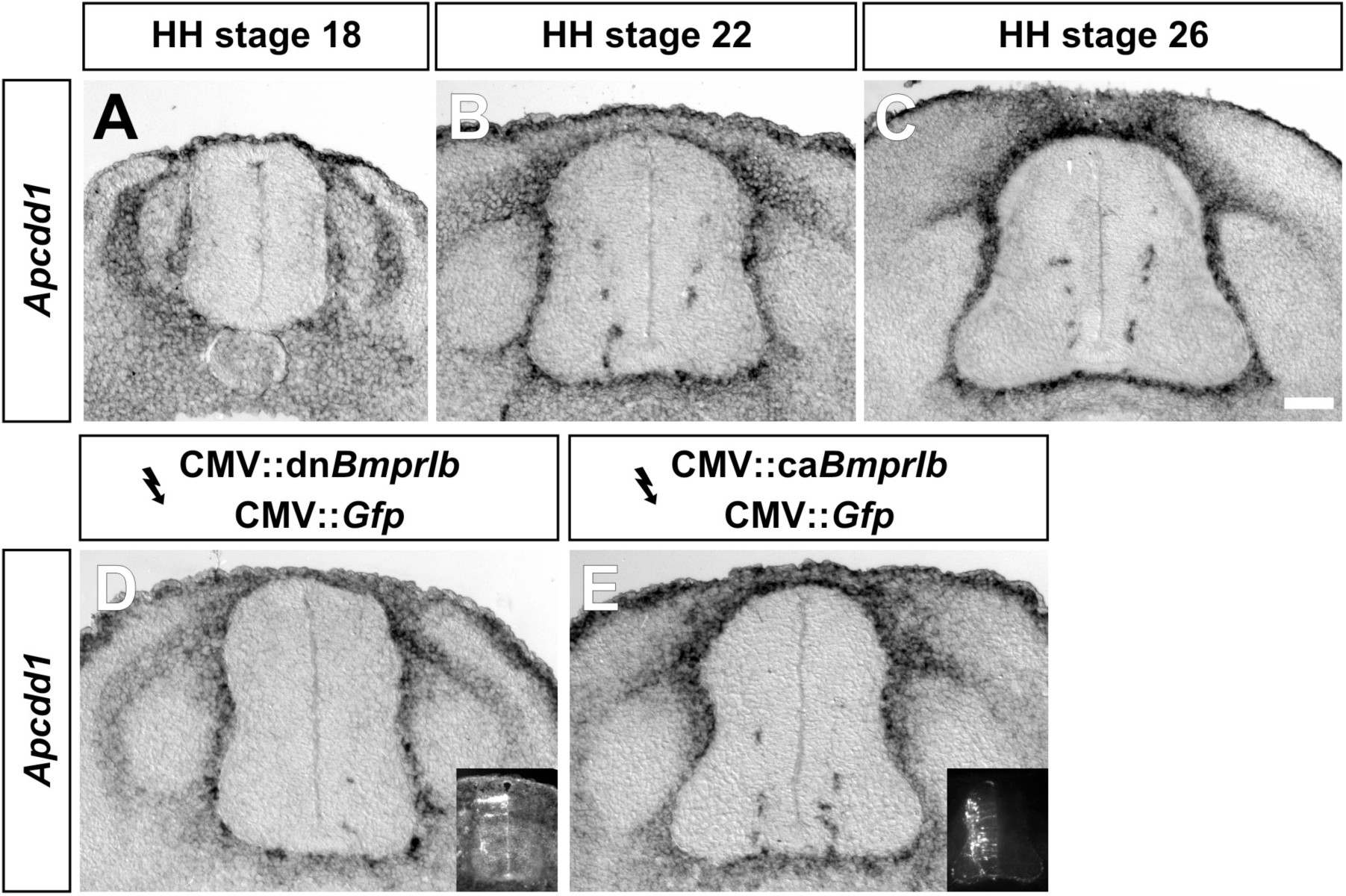
Chicken expression of *Apcdd1* in the spinal cord. **A-C.** *In situ* hybridization experiments for *Apcdd1* on transverse sections of the spinal cord taken from HH stages 18 (A), 22 (B) and 26 (C) chicken embryos. *Apcdd1* is expressed in the presumptive epidermis, as well as blood vessels within the spinal cord and a population of cells surrounding the spinal cord that have a distribution similar to migrating neural crest cells. (D, E) There is no alteration of *Apdcc1* expression in the spinal cord after electroporation with either dominant negative (dn, D) or constitutively active (ca, E) *BmprIb*. Scale bar: A: 50 μm, B-E: 100 μm.

**Figure S6.**
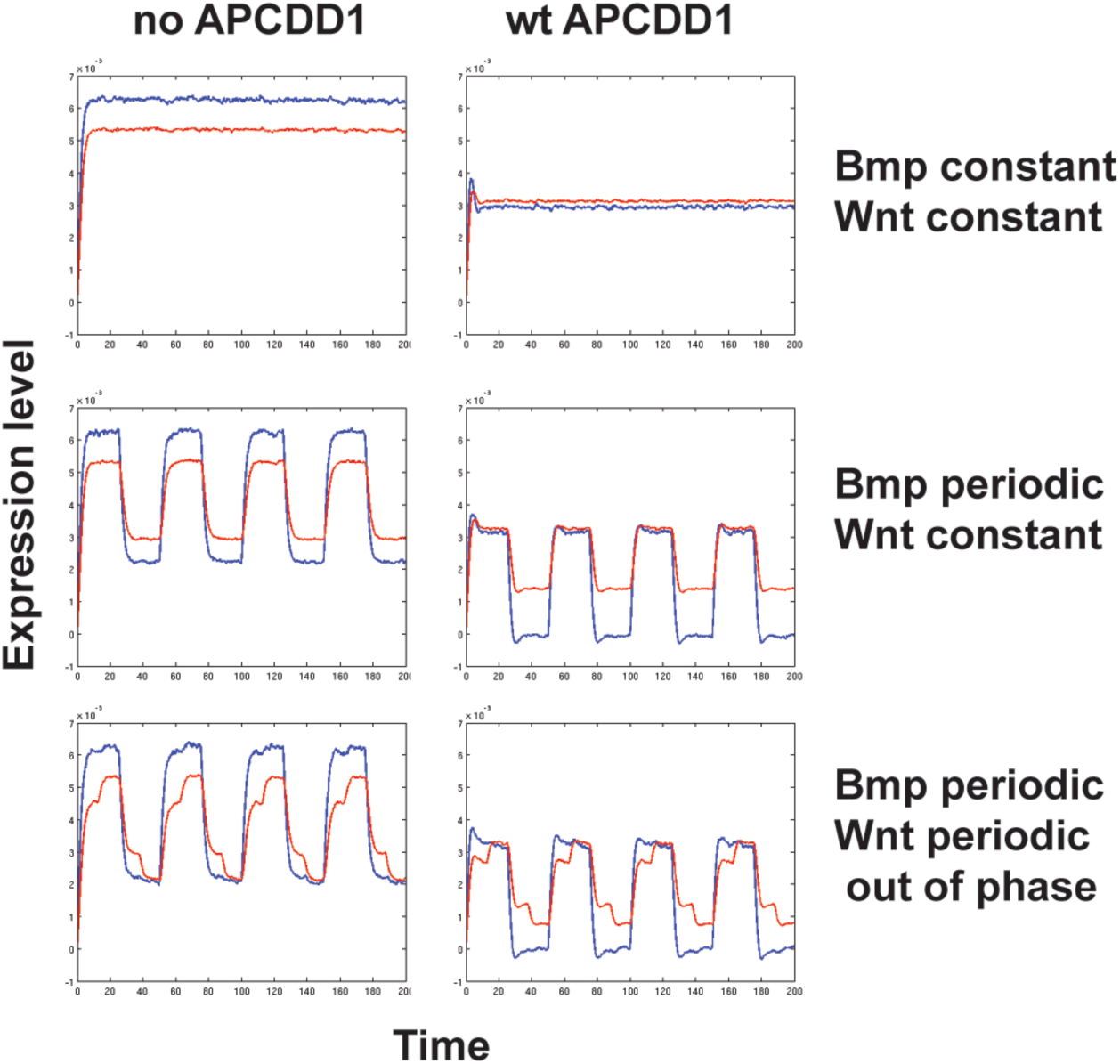
Apcdd1 may coordinate pathway dynamics and is present in multiple animal phyla. Mathematical model of the temporal activation dynamics of the BMP and Wnt signaling pathways in NIH 3T3 cells. Cells were cultured in the presence of BMP and /or Wnt (as shown in Fig. 1A-C). We modeled the levels of BMP pathway effector pSmad1 (blue) and Wnt pathway effector β-Catenin (red) against time. Each row shows one case, and each case has two subcases: no APCDD1 is present (left), Apcdd1 is present (right). Top row: when BMP ligand input and Wnt ligand input are both constant, results are consistent with observations (Fig. 1A-C). Middle row: if BMP is constant and Wnt is periodic, the model predicts both pathway effectors become periodic. Bottom row (middle row): if BMP and Wnt are both periodic and out of phase by ¼ of a period, the model predicts a complex periodic dynamics of effectors. In all three cases, the presence of Apcdd1 (right column) lowers the levels of both pSmad1 and β-Catenin compared to when it is absent (left column).

**Figure S7.**
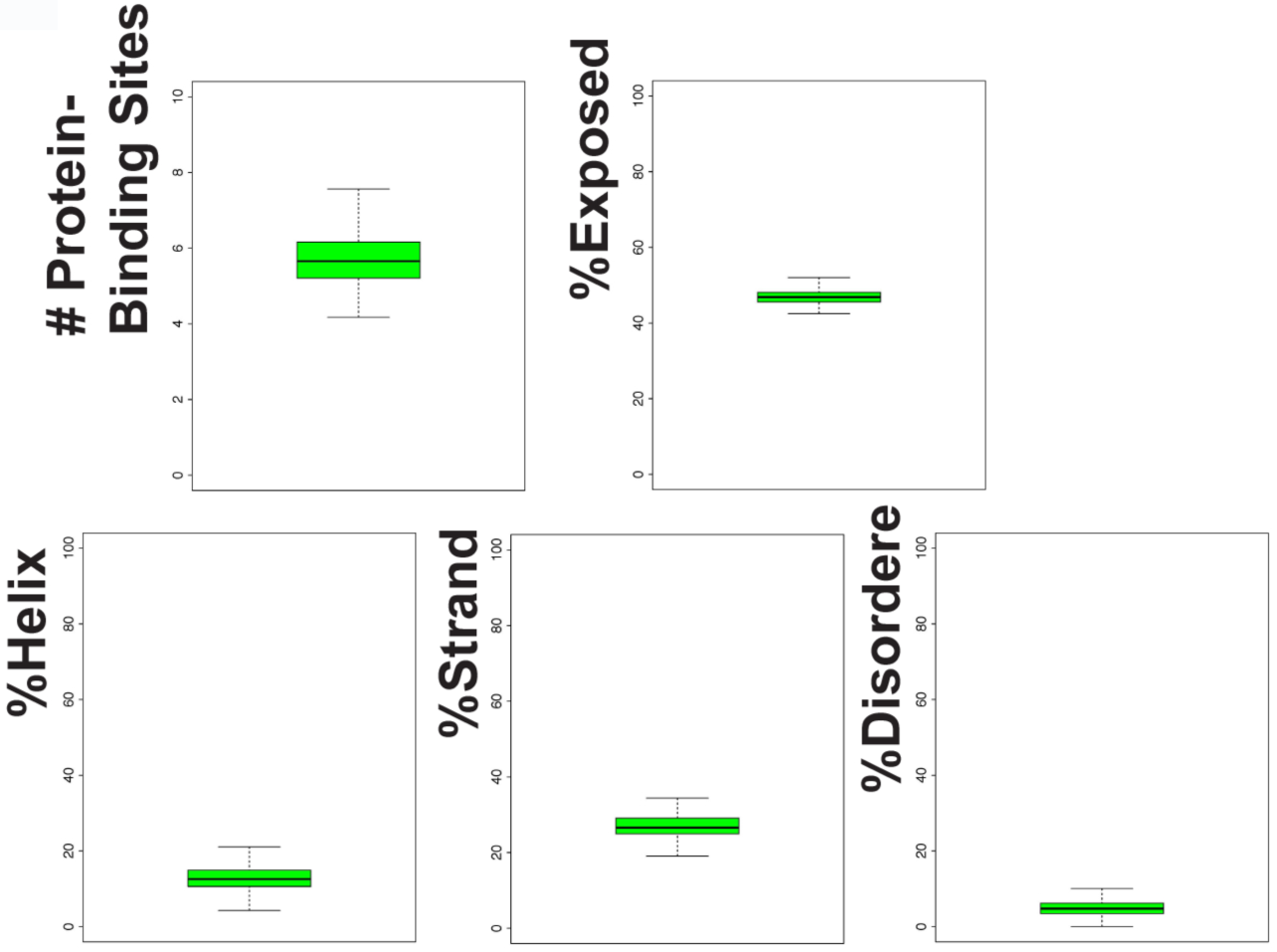
Protein structure analysis of Apcdd1 proteins. Secondary structure analysis of 441 Apcdd1 proteins shows the presence of alpha-helices, beta-strands, and protein-binding sites. Apcdd1 proteins have low disorder and low exposure to water.

## References

Abrusan, G., 2013. Integration of new genes into cellular networks, and their structural maturation. Genetics 195, 1407–1417.

Altschul, S.F., Gish, W., Miller, W., Myers, E.W., Lipman, D.J., 1990. Basic local alignment search tool. J Mol Biol 215, 403–410.

Angerer, L.M., Oleksyn, D.W., Logan, C.Y., McClay, D.R., Dale, L., Angerer, R.C., 2000. A BMP pathway regulates cell fate allocation along the sea urchin animal-vegetal embryonic axis. Development 127, 1105–1114.

Augsburger, A., Schuchardt, A., Hoskins, S., Dodd, J., Butler, S., 1999. BMPs as mediators of roof plate repulsion of commissural neurons. Neuron 24, 127–141.

Biegert, A., Soding, J., 2008. De novo identification of highly diverged protein repeats by probabilistic consistency. Bioinformatics 24, 807–814.

Bijakowski, C., Vadon-Le Goff, S., Delolme, F., Bourhis, J.M., Lecorche, P., Ruggiero, F., Becker-Pauly, C., Yiallouros, I., Stocker, W., Dive, V., Hulmes, D.J., Moali, C., 2012. Sizzled is unique among secreted frizzled-related proteins for its ability to specifically inhibit bone morphogenetic protein-1 (BMP-1)/tolloid-like proteinases. J Biol Chem 287, 33581–33593.

Bradley, L., Sun, B., Collins-Racie, L., LaVallie, E., McCoy, J., Sive, H., 2000. Different activities of the frizzled-related proteins frzb2 and sizzled2 during Xenopus anteroposterior patterning. Dev Biol 227, 118–132.

Briscoe, J., Pierani, A., Jessell, T.M., Ericson, J., 2000. A homeodomain protein code specifies progenitor cell identity and neuronal fate in the ventral neural tube. Cell 101, 435–445.

Bu, Q., Li, Z., Zhang, J., Xu, F., Liu, J., Liu, H., 2017. The crystal structure of full-length Sizzled from Xenopus laevis yields insights into Wnt-antagonistic function of secreted Frizzled-related proteins. J Biol Chem 292, 16055–16069.

Carvunis, A.R., Rolland, T., Wapinski, I., Calderwood, M.A., Yildirim, M.A., Simonis, N., Charloteaux, B., Hidalgo, C.A., Barbette, J., Santhanam, B., Brar, G.A., Weissman, J.S., Regev, A., Thierry-Mieg, N., Cusick, M.E., Vidal, M., 2012. Proto-genes and de novo gene birth. Nature 487, 370–374.

Chen, Y., Zhang, Y., Tang, J., Liu, F., Hu, Q., Luo, C., Tang, J., Feng, H., Zhang, J.H., 2015. Norrin protected blood-brain barrier via frizzled-4/beta-catenin pathway after subarachnoid hemorrhage in rats. Stroke 46, 529–536.

Chizhikov, V.V., Millen, K.J., 2005. Roof plate-dependent patterning of the vertebrate dorsal central nervous system. Dev Biol 277, 287–295.

Crooks, G.E., Hon, G., Chandonia, J.M., Brenner, S.E., 2004. WebLogo: a sequence logo generator. Genome Res 14, 1188–1190.

Cruciat, C.M., Niehrs, C., 2013. Secreted and transmembrane wnt inhibitors and activators. Cold Spring Harb Perspect Biol 5, a015081.

Daneman, R., Zhou, L., Kebede, A.A., Barres, B.A., 2010. Pericytes are required for blood-brain barrier integrity during embryogenesis. Nature 468, 562–566.

Darras, S., Fritzenwanker, J.H., Uhlinger, K.R., Farrelly, E., Pani, A.M., Hurley, I.A., Norris, R.P., Osovitz, M., Terasaki, M., Wu, M., Aronowicz, J., Kirschner, M., Gerhart, J.C., Lowe, C.J., 2018. Anteroposterior axis patterning by early canonical Wnt signaling during hemichordate development. PLoS Biol 16, e2003698.

Eddy, S.R., 1998. Profile hidden Markov models. Bioinformatics 14, 755–763.

Edgar, R.C., 2004. MUSCLE: multiple sequence alignment with high accuracy and high throughput. Nucleic Acids Res 32, 1792–1797.

Edwards, S.V., Liu, L., Pearl, D.K., 2007. High-resolution species trees without concatenation. Proc Natl Acad Sci U S A 104, 5936–5941.

El-Gebali, S., Mistry, J., Bateman, A., Eddy, S.R., Luciani, A., Potter, S.C., Qureshi, M., Richardson, L.J., Salazar, G.A., Smart, A., Sonnhammer, E.L.L., Hirsh, L., Paladin, L., Piovesan, D., Tosatto, S.C.E., Finn, R.D., 2019. The Pfam protein families database in 2019. Nucleic Acids Res 47, D427–D432.

Guder, C., Philipp, I., Lengfeld, T., Watanabe, H., Hobmayer, B., Holstein, T.W., 2006. The Wnt code: cnidarians signal the way. Oncogene 25, 7450–7460.

Hamburger, V., Hamilton, H.L., 1992. A series of normal stages in the development of the chick embryo. 1951. Dev Dyn 195, 231–272.

Hartnett, L., Glynn, C., Nolan, C.M., Grealy, M., Byrnes, L., 2010. Insulin-like growth factor-2 regulates early neural and cardiovascular system development in zebrafish embryos. Int J Dev Biol 54, 573–583.

Hata, A., Seoane, J., Lagna, G., Montalvo, E., Hemmati-Brivanlou, A., Massague, J., 2000. OAZ uses distinct DNA- and protein-binding zinc fingers in separate BMP-Smad and Olf signaling pathways. Cell 100, 229–240.

Hazen, V.M., Andrews, M.A., Umans, L., Crenshaw, E.B., 3rd, Zwijsen, A., Butler, S.J., 2012. BMP receptor-activated Smads direct diverse functions during the development of the dorsal spinal cord. Dev Biol 367, 216–227.

Hazen, V.M., Phan, K.D., Hudiburgh, S., Butler, S.J., 2011. Inhibitory Smads differentially regulate cell fate specification and axon dynamics in the dorsal spinal cord. Dev Biol 356, 566–575.

Henry, J.Q., Perry, K.J., Martindale, M.Q., 2010. beta-catenin and early development in the gastropod, Crepidula fornicata. Integr Comp Biol 50, 707–719.

Henry, J.Q., Perry, K.J., Wever, J., Seaver, E., Martindale, M.Q., 2008. Beta-catenin is required for the establishment of vegetal embryonic fates in the nemertean, Cerebratulus lacteus. Dev Biol 317, 368–379.

Hikasa, H., Sokol, S.Y., 2013. Wnt signaling in vertebrate axis specification. Cold Spring Harb Perspect Biol 5, a007955.

Hogvall, M., Schonauer, A., Budd, G.E., McGregor, A.P., Posnien, N., Janssen, R., 2014. Analysis of the Wnt gene repertoire in an onychophoran provides new insights into the evolution of segmentation. Evodevo 5, 14.

Hsiao, C.D., You, M.S., Guh, Y.J., Ma, M., Jiang, Y.J., Hwang, P.P., 2007. A positive regulatory loop between foxi3a and foxi3b is essential for specification and differentiation of zebrafish epidermal ionocytes. PLoS One 2, e302.

Jukkola, T., Sinjushina, N., Partanen, J., 2004. Drapc1 expression during mouse embryonic development. Gene Expr Patterns 4, 755–762.

Kandyba, E., Leung, Y., Chen, Y.B., Widelitz, R., Chuong, C.M., Kobielak, K., 2013. Competitive balance of intrabulge BMP/Wnt signaling reveals a robust gene network ruling stem cell homeostasis and cyclic activation. Proc Natl Acad Sci U S A 110, 1351–1356.

Kao, K.R., Elinson, R.P., 1988. The entire mesodermal mantle behaves as Spemann’s organizer in dorsoanterior enhanced Xenopus laevis embryos. Dev Biol 127, 64–77.

Keener, J., Sneyd, J., 2009. Mathematical Physiology I: Cellular Physiology. Springer, New York.

Korchynskyi, O., ten Dijke, P., 2002. Identification and functional characterization of distinct critically important bone morphogenetic protein-specific response elements in the Id1 promoter. J Biol Chem 277, 4883–4891.

Kumar, S., Stecher, G., Suleski, M., Hedges, S.B., 2017. TimeTree: A Resource for Timelines, Timetrees, and Divergence Times. Mol Biol Evol 34, 1812–1819.

Kuo, D.H., Weisblat, D.A., 2011. A new molecular logic for BMP-mediated dorsoventral patterning in the leech Helobdella. Curr Biol 21, 1282–1288.

Kuraku, S., Usuda, R., Kuratani, S., 2005. Comprehensive survey of carapacial ridge-specific genes in turtle implies co-option of some regulatory genes in carapace evolution. Evol Dev 7, 3–17.

Kwon, H.J., Bhat, N., Sweet, E.M., Cornell, R.A., Riley, B.B., 2010. Identification of early requirements for preplacodal ectoderm and sensory organ development. PLoS Genet 6, e1001133.

Lambert, J.D., Johnson, A.B., Hudson, C.N., Chan, A., 2016. Dpp/BMP2-4 Mediates Signaling from the D-Quadrant Organizer in a Spiralian Embryo. Curr Biol 26, 2003–2010.

Lanfear, R., Frandsen, P.B., Wright, A.M., Senfeld, T., Calcott, B., 2017. PartitionFinder 2: New Methods for Selecting Partitioned Models of Evolution for Molecular and Morphological Phylogenetic Analyses. Mol Biol Evol 34, 772–773.

Lee, H.X., Ambrosio, A.L., Reversade, B., De Robertis, E.M., 2006. Embryonic dorsal-ventral signaling: secreted frizzled-related proteins as inhibitors of tolloid proteinases. Cell 124, 147–159.

Lee, J., Tumbar, T., 2012. Hairy tale of signaling in hair follicle development and cycling. Semin Cell Dev Biol 23, 906–916.

Lee, K.J., Dietrich, P., Jessell, T.M., 2000. Genetic ablation reveals that the roof plate is essential for dorsal interneuron specification. Nature 403, 734–740.

Liem, K.F., Jr., Tremml, G., Jessell, T.M., 1997. A role for the roof plate and its resident TGFbeta-related proteins in neuronal patterning in the dorsal spinal cord. Cell 91, 127–138.

Lim, X., Nusse, R., 2013. Wnt signaling in skin development, homeostasis, and disease. Cold Spring Harb Perspect Biol 5.

Lin, L.Y., Horng, J.L., Kunkel, J.G., Hwang, P.P., 2006. Proton pump-rich cell secretes acid in skin of zebrafish larvae. Am J Physiol Cell Physiol 290, C371–378.

Lowe, C.J., Terasaki, M., Wu, M., Freeman, R.M., Jr., Runft, L., Kwan, K., Haigo, S., Aronowicz, J., Lander, E., Gruber, C., Smith, M., Kirschner, M., Gerhart, J., 2006. Dorsoventral patterning in hemichordates: insights into early chordate evolution. PLoS Biol 4, e291.

Lu, F.I., Thisse, C., Thisse, B., 2011. Identification and mechanism of regulation of the zebrafish dorsal determinant. Proc Natl Acad Sci U S A 108, 15876–15880.

Marlow, H., Tosches, M.A., Tomer, R., Steinmetz, P.R., Lauri, A., Larsson, T., Arendt, D., 2014. Larval body patterning and apical organs are conserved in animal evolution. BMC Biol 12, 7.

Mazzoni, J., Smith, J.R., Shahriar, S., Cutforth, T., Ceja, B., Agalliu, D., 2017. The Wnt Inhibitor Apcdd1 Coordinates Vascular Remodeling and Barrier Maturation of Retinal Blood Vessels. Neuron 96, 1055–1069 e1056.

Miyoshi, H., Ajima, R., Luo, C.T., Yamaguchi, T.P., Stappenbeck, T.S., 2012. Wnt5a potentiates TGF-beta signaling to promote colonic crypt regeneration after tissue injury. Science 338, 108–113.

Mizutani, C.M., Bier, E., 2008. EvoD/Vo: the origins of BMP signalling in the neuroectoderm. Nat Rev Genet 9, 663–677.

Molina, M.D., Salo, E., Cebria, F., 2011. Organizing the DV axis during planarian regeneration. Commun Integr Biol 4, 498–500.

Moustakas, A., Heldin, C.H., 2009. The regulation of TGFbeta signal transduction. Development 136, 3699–3714.

Mueller, T.D., Nickel, J., 2012. Promiscuity and specificity in BMP receptor activation. FEBS Lett 586, 1846–1859.

Muller-Rover, S., Handjiski, B., van der Veen, C., Eichmuller, S., Foitzik, K., McKay, I.A., Stenn, K.S., Paus, R., 2001. A comprehensive guide for the accurate classification of murine hair follicles in distinct hair cycle stages. J Invest Dermatol 117, 3–15.

Mulloy, B., Rider, C.C., 2015. The Bone Morphogenetic Proteins and Their Antagonists. Vitam Horm 99, 63–90.

Muraoka, O., Shimizu, T., Yabe, T., Nojima, H., Bae, Y.K., Hashimoto, H., Hibi, M., 2006. Sizzled controls dorso-ventral polarity by repressing cleavage of the Chordin protein. Nat Cell Biol 8, 329–338.

Nusslein-Volhard, C., Wieschaus, E., 1980. Mutations affecting segment number and polarity in Drosophila. Nature 287, 795–801.

Oz-Levi, D., Olender, T., Bar-Joseph, I., Zhu, Y., Marek-Yagel, D., Barozzi, I., Osterwalder, M., Alkelai, A., Ruzzo, E.K., Han, Y., Vos, E.S.M., Reznik-Wolf, H., Hartman, C., Shamir, R., Weiss, B., Shapiro, R., Pode-Shakked, B., Tatarskyy, P., Milgrom, R., Schvimer, M., Barshack, I., Imai, D.M., Coleman-Derr, D., Dickel, D.E., Nord, A.S., Afzal, V., van Bueren, K.L., Barnes, R.M., Black, B.L., Mayhew, C.N., Kuhar, M.F., Pitstick, A., Tekman, M., Stanescu, H.C., Wells, J.M., Kleta, R., de Laat, W., Goldstein, D.B., Pras, E., Visel, A., Lancet, D., Anikster, Y., Pennacchio, L.A., 2019. Noncoding deletions reveal a gene that is critical for intestinal function. Nature 571, 107–111.

Pang, K., Ryan, J.F., Baxevanis, A.D., Martindale, M.Q., 2011. Evolution of the TGF-beta signaling pathway and its potential role in the ctenophore, Mnemiopsis leidyi. PLoS One 6, e24152.

Pang, K., Ryan, J.F., Program, N.C.S., Mullikin, J.C., Baxevanis, A.D., Martindale, M.Q., 2010. Genomic insights into Wnt signaling in an early diverging metazoan, the ctenophore Mnemiopsis leidyi. Evodevo 1, 10.

Park, F.D., Priess, J.R., 2003. Establishment of POP-1 asymmetry in early C. elegans embryos. Development 130, 3547–3556.

Phan, K.D., Hazen, V.M., Frendo, M., Jia, Z., Butler, S.J., 2010. The bone morphogenetic protein roof plate chemorepellent regulates the rate of commissural axonal growth. J Neurosci 30, 15430–15440.

Piccolo, S., Agius, E., Leyns, L., Bhattacharyya, S., Grunz, H., Bouwmeester, T., De Robertis, E.M., 1999. The head inducer Cerberus is a multifunctional antagonist of Nodal, BMP and Wnt signals. Nature 397, 707–710.

Pires-daSilva, A., Sommer, R.J., 2003. The evolution of signalling pathways in animal development. Nat Rev Genet 4, 39–49.

Plickert, G., Jacoby, V., Frank, U., Muller, W.A., Mokady, O., 2006. Wnt signaling in hydroid development: formation of the primary body axis in embryogenesis and its subsequent patterning. Dev Biol 298, 368–378.

Plikus, M.V., Baker, R.E., Chen, C.C., Fare, C., de la Cruz, D., Andl, T., Maini, P.K., Millar, S.E., Widelitz, R., Chuong, C.M., 2011. Self-organizing and stochastic behaviors during the regeneration of hair stem cells. Science 332, 586–589.

Plikus, M.V., Mayer, J.A., de la Cruz, D., Baker, R.E., Maini, P.K., Maxson, R., Chuong, C.M., 2008. Cyclic dermal BMP signalling regulates stem cell activation during hair regeneration. Nature 451, 340–344.

Plouhinec, J.L., Zakin, L., De Robertis, E.M., 2011. Systems control of BMP morphogen flow in vertebrate embryos. Curr Opin Genet Dev 21, 696–703.

Rentzsch, F., Guder, C., Vocke, D., Hobmayer, B., Holstein, T.W., 2007. An ancient chordin-like gene in organizer formation of Hydra. Proc Natl Acad Sci U S A 104, 3249–3254.

Reversade, B., De Robertis, E.M., 2005. Regulation of ADMP and BMP2/4/7 at opposite embryonic poles generates a self-regulating morphogenetic field. Cell 123, 1147–1160.

Reversade, B., Kuroda, H., Lee, H., Mays, A., De Robertis, E.M., 2005. Depletion of Bmp2, Bmp4, Bmp7 and Spemann organizer signals induces massive brain formation in Xenopus embryos. Development 132, 3381–3392.

Salic, A.N., Kroll, K.L., Evans, L.M., Kirschner, M.W., 1997. Sizzled: a secreted Xwnt8 antagonist expressed in the ventral marginal zone of Xenopus embryos. Development 124, 4739–4748.

Sasai, Y., Lu, B., Steinbeisser, H., De Robertis, E.M., 1995. Regulation of neural induction by the Chd and Bmp-4 antagonistic patterning signals in Xenopus. Nature 376, 333–336.

Schaefer, C., Schlessinger, A., Rost, B., 2010. Protein secondary structure appears to be robust under in silico evolution while protein disorder appears not to be. Bioinformatics 26, 625–631.

Schenkelaars, Q., Pratlong, M., Kodjabachian, L., Fierro-Constain, L., Vacelet, J., Le Bivic, A., Renard, E., Borchiellini, C., 2017. Animal multicellularity and polarity without Wnt signaling. Sci Rep 7, 15383.

Scimone, M.L., Cote, L.E., Rogers, T., Reddien, P.W., 2016. Two FGFRL-Wnt circuits organize the planarian anteroposterior axis. Elife 5.

Shimomura, Y., Agalliu, D., Vonica, A., Luria, V., Wajid, M., Baumer, A., Belli, S., Petukhova, L., Schinzel, A., Brivanlou, A.H., Barres, B.A., Christiano, A.M., 2010. APCDD1 is a novel Wnt inhibitor mutated in hereditary hypotrichosis simplex. Nature 464, 1043–1047.

Song, J., McColl, J., Camp, E., Kennerley, N., Mok, G.F., McCormick, D., Grocott, T., Wheeler, G.N., Munsterberg, A.E., 2014. Smad1 transcription factor integrates BMP2 and Wnt3a signals in migrating cardiac progenitor cells. Proc Natl Acad Sci U S A 111, 7337–7342.

Soza-Ried, C., Hess, I., Netuschil, N., Schorpp, M., Boehm, T., 2010. Essential role of c-myb in definitive hematopoiesis is evolutionarily conserved. Proc Natl Acad Sci U S A 107, 17304–17308.

Stamatakis, A., 2014. RAxML version 8: a tool for phylogenetic analysis and post-analysis of large phylogenies. Bioinformatics 30, 1312–1313.

Suel, G.M., Kulkarni, R.P., Dworkin, J., Garcia-Ojalvo, J., Elowitz, M.B., 2007. Tunability and noise dependence in differentiation dynamics. Science 315, 1716–1719.

Suzuki, Y., Yandell, M.D., Roy, P.J., Krishna, S., Savage-Dunn, C., Ross, R.M., Padgett, R.W., Wood, W.B., 1999. A BMP homolog acts as a dose-dependent regulator of body size and male tail patterning in Caenorhabditis elegans. Development 126, 241–250.

Tamura, K., Peterson, D., Peterson, N., Stecher, G., Nei, M., Kumar, S., 2011. MEGA5: molecular evolutionary genetics analysis using maximum likelihood, evolutionary distance, and maximum parsimony methods. Mol Biol Evol 28, 2731–2739.

Tautz, D., Domazet-Loso, T., 2011. The evolutionary origin of orphan genes. Nat Rev Genet 12, 692–702.

Timmer, J.R., Wang, C., Niswander, L., 2002. BMP signaling patterns the dorsal and intermediate neural tube via regulation of homeobox and helix-loop-helix transcription factors. Development 129, 2459–2472.

Treffkorn, S., Mayer, G., 2013. Expression of the decapentaplegic ortholog in embryos of the onychophoran Euperipatoides rowelli. Gene Expr Patterns 13, 384–394.

Tsuchida, T., Ensini, M., Morton, S.B., Baldassare, M., Edlund, T., Jessell, T.M., Pfaff, S.L., 1994. Topographic organization of embryonic motor neurons defined by expression of LIM homeobox genes. Cell 79, 957–970.

Umulis, D., O’Connor, M.B., Blair, S.S., 2009. The extracellular regulation of bone morphogenetic protein signaling. Development 136, 3715–3728.

Vonica, A., Rosa, A., Arduini, B.L., Brivanlou, A.H., 2011. APOBEC2, a selective inhibitor of TGFbeta signaling, regulates left-right axis specification during early embryogenesis. Dev Biol 350, 13–23.

Wikramanayake, A.H., Huang, L., Klein, W.H., 1998. beta-Catenin is essential for patterning the maternally specified animal-vegetal axis in the sea urchin embryo. Proc Natl Acad Sci U S A 95, 9343–9348.

Windsor, P.J., Leys, S.P., 2010. Wnt signaling and induction in the sponge aquiferous system: evidence for an ancient origin of the organizer. Evol Dev 12, 484–493.

Wine-Lee, L., Ahn, K.J., Richardson, R.D., Mishina, Y., Lyons, K.M., Crenshaw, E.B., 3rd, 2004. Signaling through BMP type 1 receptors is required for development of interneuron cell types in the dorsal spinal cord. Development 131, 5393–5403.

Xu, B., Chen, C., Chen, H., Zheng, S.G., Bringas, P., Jr., Xu, M., Zhou, X., Chen, D., Umans, L., Zwijsen, A., Shi, W., 2011. Smad1 and its target gene Wif1 coordinate BMP and Wnt signaling activities to regulate fetal lung development. Development 138, 925–935.

Yabe, T., Shimizu, T., Muraoka, O., Bae, Y.K., Hirata, T., Nojima, H., Kawakami, A., Hirano, T., Hibi, M., 2003. Ogon/Secreted Frizzled functions as a negative feedback regulator of Bmp signaling. Development 130, 2705–2716.

Yachdav, G., Kloppmann, E., Kajan, L., Hecht, M., Goldberg, T., Hamp, T., Honigschmid, P., Schafferhans, A., Roos, M., Bernhofer, M., Richter, L., Ashkenazy, H., Punta, M., Schlessinger, A., Bromberg, Y., Schneider, R., Vriend, G., Sander, C., Ben-Tal, N., Rost, B., 2014. PredictProtein--an open resource for online prediction of protein structural and functional features. Nucleic Acids Res 42, W337–343.

Yamauchi, K., Varadarajan, S.G., Li, J.E., Butler, S.J., 2013. Type Ib BMP receptors mediate the rate of commissural axon extension through inhibition of cofilin activity. Development 140, 333–342.

Yu, P.B., Hong, C.C., Sachidanandan, C., Babitt, J.L., Deng, D.Y., Hoyng, S.A., Lin, H.Y., Bloch, K.D., Peterson, R.T., 2008. Dorsomorphin inhibits BMP signals required for embryogenesis and iron metabolism. Nat Chem Biol 4, 33–41.

