## Supplemental Table 1 for "Apcdd1 is a dual BMP/Wnt inhibitor in the developing nervous system and skin"

| Probe Set ID | Fold change ([Depl] vs [Control]) | Regulation ([Depl] vs [Control]) | [Control](raw) | [Depl](raw) | Gene Symbol | Transcript Assignments |
|---|---|---|---|---|---|---|
| **UPREGULATED** | | | | | | |
| Xl2.20089.1.S2_at | 13.29261 | up | 54.168644 | 717.92413 | foxi1-ema/Xema/Foxi1/HNF-3 | BC042303 //forkhead box I1, mRNA (cDNA clone MGC:52689 IMAGE:4682602). |
| Xl2.54072.1.S1_at | 10.586516 | up | 241.02812 | 2517.7058 | H3r | DQ096858 // h3r mRNA. |
| Xl2.12115.1.S1_s_at | 8.925359 | up | 42.059174 | 371.29764 | plk2 | AF357840 // s polo-like kinase 2 (Plx2) mRNA. |
| Xl2.55726.1.A1_at | 6.2248645 | up | 57.024197 | 356.19614 | ccng1 | BC126020 // cyclin G1, mRNA (cDNA clone MGC:154808 IMAGE:8328440). |
| Xl2.49984.1.S1_at | 4.865988 | up | 10.59639 | 51.290554 | PARP3 | NM_001095376 // hypothetical LOC496154 (LOC496154), mRNA. |
| Xl2.13019.1.S1_at | 3.4627233 | up | 483.91782 | 1676.7993 | rasl11b | NM_001085926 // hypothetical protein MGC53580 (MGC53580), mRNA. |
| Xl2.21079.2.S1_a_at | 3.189228 | up | 1308.8922 | 4149.4937 | h2B | BC077399 // histone H2B, mRNA (cDNA clone MGC:81709 IMAGE:6865010). |
| Xl2.49747.1.S1_at | 2.9899995 | up | 13.932175 | 41.671738 | cytosolic 5'-nucleotidase III | NM_001094563 // hypothetical LOC494724 (LOC494724), mRNA. |
| Xl2.23649.1.S1_at | 2.806288 | up | 374.38474 | 1051.3436 | agtrl1 | BC139486 // angiotensin receptor related protein, mRNA (cDNA clone MGC:154551 IMAGE:8321886). |
| Xl2.23887.1.S2_at | 2.4871519 | up | 86.91137 | 216.20465 | RAB40B | BC078543 // /DEF= MGC85390 protein, mRNA (cDNA clone MGC:85390 IMAGE:6931765). |
| Xl2.16135.1.A1_at | 2.4030888 | up | 61.00638 | 146.3802 | Fgfr2 | BJ045607 // /DB_XREF=BJ045607 /CLONE=XL004l08 |
| Xl2.1475.1.A1_at | 2.2238114 | up | 152.57167 | 337.7993 | pik3r2 | BC077814 // phosphoinositide-3-kinase, regulatory subunit, polypeptide 2 (p85 beta), mRNA (cDNA clone MGC:80449 IMAGE:5155911). |
| Xl2.6970.1.S1_at | 2.2069082 | up | 83.7789 | 185.60034 | NADH DH (ubiquinone) 1 alpha subcomplex, 8, 19kDa | NM_001096329 // hypothetical protein MGC131357 (MGC131357), mRNA. |
| Xl2.8458.1.S1_at | 2.1906917 | up | 210.4199 | 462.8243 | E3 ubq-protein ligase MARCH8 | BC084236 // E3 ubiquitin-protein ligase MARCH8, mRNA (cDNA clone MGC:81586 IMAGE:6863461). |
| Xl2.24839.1.S2_s_at | 2.1296399 | up | 21.284641 | 45.323395 | mafB | BC077255 // bZIP transcription factor mafB, mRNA (cDNA clone MGC:79895 IMAGE:4965723). |
| Xl2.49806.1.S1_at | 2.0961912 | up | 287.9446 | 603.6631 | ndufa10b | BC068897 // NADH dehydrogenase (ubiquinone) 1 alpha subcomplex, 10, 42kDa, mRNA (cDNA clone MGC:83091 IMAGE:6317233). |
| Xl2.3027.1.S1_at | 2.0226674 | up | 113.94761 | 230.9705 | psmd14/POH1/rpn11 | BC045094 // 26S proteasome-associated pad1 homolog, mRNA (cDNA clone MGC:53911 IMAGE:5569802. |

**DOWNREGULATED**

| | | | | | | |
|---|---|---|---|---|---|---|
| Xl2.16206.1.S1_at | 3.2757027 | down | 324.993 | 98.9578 | pnp | NM_001086340 // hypothetical protein MGC64592 (MGC64592), mRNA. |
| Xl2.971.1.S1_at | 3.273429 | down | 56.62768 | 17.340136 | ATP4A | NM_001090874 // ATPase, H+/K+ exchanging, alpha polypeptide (ATP4A), mRNA. |
| Xl2.32186.1.S1_at | 3.2680411 | down | 213.41772 | 65.236115 | Tyr aminotransferase | BC074414 // hypothetical protein LOC443707, mRNA (cDNA clone IMAGE:7019538), partial cds. |
| Xl2.55911.1.S1_at | 2.9581723 | down | 338.9496 | 114.06936 | FPGS | BI477984 // /DB_XREF=dae60g04.y3 /CLONE=IMAGE:4678471 |
| Xl2.23438.1.S3_at | 2.7084556 | down | 330.64606 | 121.7796 | hnrnpu | BC046700 // /DEF=, Similar to heterogeneous nuclear ribonucleoprotein U (scaffold attachment factor A), clone IMAGE:5542936, mRNA, partial cds. |
| Xl2.48751.1.S1_at | 2.673174 | down | 324.07248 | 121.10416 | eif1ad | NM_001095509 // hypothetical LOC496359 (eif1ad), mRNA. |
| Xl2.706.1.S1_at | 2.6462636 | down | 80.22797 | 30.232681 | blm-A | AF307841 // Bloom's syndrome-like protein mRNA, complete cds. |
| Xl2.24047.1.S1_at | 2.542717 | down | 181.5669 | 71.42785 | slc38a2/SNAT2 | BC077990 // solute carrier family 38, member 2, mRNA (cDNA clone MGC:81747 IMAGE:6865688), complete cds. |
| Xl2.47730.1.S1_at | 2.325578 | down | 286.89365 | 123.314354 | rdh 10-A | BC077913 // MGC80820 protein, mRNA (cDNA clone MGC:80820 IMAGE:5513889). |
| Xl2.47155.1.S1_at | 2.2501059 | down | 151.47588 | 67.67626 | ttc5 | BC126001 // hypothetical protein LOC431931, mRNA (cDNA clone MGC:154647 IMAGE:8330240). |
| Xl2.7874.1.A1_at | 2.2132833 | down | 202.09216 | 91.53327 | plekhm3 | BC124982 // Pleckstrin homology domain-containing family M member 3, mRNA (cDNA clone MGC:154846 IMAGE:8330058). |
| Xl2.1179.1.S1_at | 2.208371 | down | 207.97151 | 94.04665 | fgf3-A | BC106382 // cDNA clone MGC:130954 IMAGE:7973902, complete cds. // gb // 14 // --- |
| Xl2.50498.1.S2_at | 2.195632 | down | 213.60956 | 97.18936 | Mxi1 | NM_001095701// max interactor 1. |
| Xl2.5386.1.S1_at | 2.15772 | down | 298.2187 | 137.80058 | znf268-b | NM_001094891 // hypothetical LOC495204, mRNA. |
| Xl2.13977.1.S1_at | 2.086489 | down | 366.33646 | 175.48596 | SIRT1 | BC141734 // cDNA clone IMAGE:7211274.class III histone deacetylase |
| Xl2.4181.1.S1_at | 2.0080626 | down | 93.44522 | 46.58273 | xrcc5-a | BC077439 // X-ray repair complementing defective repair in Chinese hamster cells 5 (double-strand-break rejoining; Ku autoantigen, 80kDa), mRNA (cDNA clone MGC:82261 IMAGE:4032122). |
| Xl2.1242.1.S1_at | 2.0041463 | down | 386.6808 | 192.38673 | arg1 | BC043635 // arginase, mRNA (cDNA clone MGC:53709 IMAGE:4889599). |
